# Phenotypic plasticity shapes biofilm’s structure and fluid transport enhancing resilience

**DOI:** 10.1101/2023.07.20.549970

**Authors:** Abhirup Mookherjee, Nikhil Krishnan, Ching-Yao Toh, Joseph Knight, Akhilesh Kumar Verma, Aaron Smith, Carolina Tropini, Luis Ruiz Pestana, Diana Fusco

**Author notes:** These authors contributed equally to this work (Abhirup Mookherjee, Nikhil Krishnan, Ching-Yao Toh).

## Abstract

Phenotypic heterogeneity is one of the hallmarks of the biofilm lifestyle, where even isogenic populations give rise to spatially organized and phenotypically distinct subpopulations. One such pattern is generated by the ability of several biofilm-forming bacteria to switch between a flagellated and a matrix producing state. Here, using *Bacillus subtilis* as a model system, we investigate the role of this switch during biofilm development on a solid-air interface.

By comparing the matrix-flagella spatio-temporal patterns in wild-type biofilms with mixtures of flagella- and matrix-null mutants biofilms, we find that pattern formation does not require a phenotypic switch that enables individual cells to respond to the local environment, but can be explained by a completely stochastic switch coupled to a phenotype-dependent fitness landscape that selects phenotypes at the population level. Integration of experiments and physical models shows that the coexistence between flagellated and matrix-producing cells provides the population with enhanced resilience to environmental changes, by enabling cells to manipulate and harness the local morphological and transport properties within the biofilm. Our results not only reveal a new evolutionary advantage of phenotypic plasticity in biofilms, but also illustrate how the biology and ecology of these populations are intrinsically tied to their physical properties.

## Introduction

One of the hallmarks of bacterial biofilms is their ability to display multiple phenotypes even in isogenic populations [1–5], which provides enhanced resilience in the form of bet-hedging [6, 7], division of labor [8] and protection against cheaters [9, 10], to name a few. This phenotypic heterogeneity is thought to arise from a combination of stochasticity in gene expression [3, 11] and single-cell response to the diverse environmental cues that emerge in the complex biofilm architecture [2].

A well-known manifestation of phenotypic heterogeneity across bacterial species is the transition between a flagellated state, in which cells express the flagellar apparatus and surfactant molecules to promote movement and dispersal, and a matrix-producing state, in which cells secrete a viscoelastic extracellular matrix (ECM) that encases the cell and its surroundings [12–14]. In *Bacillus subtilis*, the master regulator of this switch, SpoA, is phosphorylated by multiple kinases in response to certain stress conditions, activating the gene expression cascade that leads to matrix production and repression of motility-associated genes [15, 16]. This environmental response has often been cited to explain the reproducible and coordinated gene expression pattern observed in *B. subtilis* biofilms grown on agar plates [14, 17]. However, reversible switching between flagellated and matrix-producing states has also been observed in single-cell studies under constant, homogeneous, and stress-free conditions [13, 18, 19], indicating that a transition between phenotypes can occur stochastically without any environmental trigger.

These observations raise two intriguing questions. First, is a flagellated–matrix switch that allows individual cells to respond to their environment necessary to explain the reproducible spatio-temporal gene-expression pattern observed during *B. subtilis* biofilm development? Second, is there any fitness advantage associated with maintaining a flagellated-matrix heterogeneous population even at small scales where the environment favors one phenotype over the other?

In the following, we address these questions using a combination of time-lapse fluorescence microscopy and physical modelling. First, we find that a purely stochastic flagellated-matrix switch coupled with an environment-modulated growth rate is (i) sufficient to explain the gene-expression pattern of *B. subtilis* biofilms and (ii) necessary to reproduce the relative distribution of knock-outs in mixed biofilms. Second, we find that, in contrast to smooth colonies, biofilms with a complex three-dimensional architecture display a previously unreported *escaping ability*, whereby cells trapped within the core of the biofilm can escape the surrounding population and establish at the front. Importantly, we observe that only populations capable of maintaining flagellated-matrix heterogeneity at small scales exhibit such behavior. By integrating fluorescence images of multiple biofilm combinations with corresponding physical models, we show that this heterogeneity is essential to generate and utilize a structure similar to a *vascular system* within the biofilm: matrix-producing cells are responsible for triggering the mechanical instabilities that lead to wrinkle formation and for generating pressure gradients via fluid adsorption from the agar, and flagellated cells are needed to take advantage of the system and travel outwards. This elegant yet simple interplay between phenotypic heterogeneity, mechanics, and fluid transport provides biofilms with an unexpected strategy to adapt to environmental changes by allowing resistant cells to travel to the front of the biofilm and establish, while maintaining the protection offered by the ECM for the remaining susceptible population.

## Results

### Flagellated-matrix gene expression pattern in *B. subtilis* biofilms is reproduced by mixtures of non-switching populations

The characteristic pattern of flagellated-matrix gene expression in *B. subtilis* biofilm grown on MSgg agar substrates (Methods), which is known to induce biofilm formation, exhibits first the appearance of an outer ring made of matrix producing cells where radial wrinkles develop until day 4, followed by the emergence of a flagellated ring at the outer edge of the biofilm around day 5 (Fig. 1, a-d, Supplementary Movie 1). The high reproducibility of this pattern has often been explained by a single-cell gene expression response to environmental conditions that vary in space and time during biofilm growth (e.g., nutrient availability, oxygen, etc..) [16, 17, 20]. Yet, high-resolution two-photon microscopy reveals phenotypic heterogeneity even at very small scales, where external factors are expected to be homogeneous [16]. This finding suggests that stochastic fluctuations in the expression of motility vs. matrix genes [13] can occur even under biofilm conditions, allowing the community to maintain local subpopulations of phenotypically heterogeneous cell types.

**Fig. 1:**
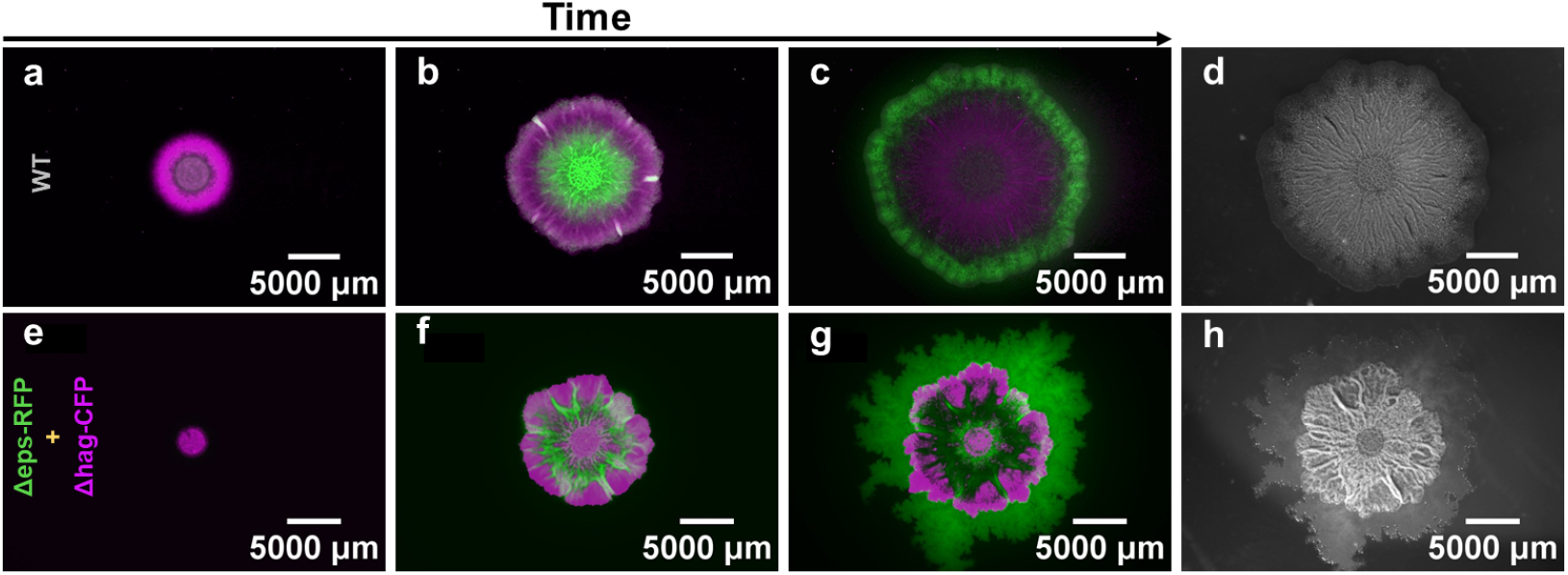
Gene expression pattern in a wild-type biofilm formed by the dual-reporter strain (a-d, magenta for matrix and green for flagella; i.e., sacA::Phag-yfp, amyE::PtapA-tsr-mCherry) and strain distribution in the mixture of flagella and matrix knock-outs (e-h, magenta for Δ*hag*-CFP and green for Δ*eps*-RFP). Panels correspond to 24 h (a), 49 h (b), 132 h (c-d), 24 h (e), 72 h (f), and 168 h (g-h).

To understand whether the presence of phenotypic heterogeneity alone is sufficient to qualitatively reproduce the gene expression pattern of the wild-type biofilm, we mixed in a single droplet two knock-out strains: Δ*hag*, lacking the main flagellar filament, and Δ*eps*, lacking a major component of the ECM, essential for wrinkle-formation [21] (Fig. 1e-g, Supplementary Movie 2). We note that in these two strains the phenotypic switch is not altered; however, each mutant exhibits impaired functions in one phenotype because of the gene knock-out. In practice, therefore, the ability of these strains to switch between functional flagellated and matrix-producing phenotypes is severely compromised, as demonstrated by the defective biofilm morphology of either strain when grown in isolation (Fig. S1).

Remarkably, we found that the spatio-temporal distribution of the mixed biofilm qualitatively mirrors the sequence of gene expression patterns observed in the wild-type (Fig. 1), with the matrix-only Δ*hag* forming a first outer ring, and the flagellated-only Δ*eps* taking over the biofilm front at later times. Importantly, the spatial distribution of the strains showed a high degree of polar symmetry, in contrast to the typical sectoring pattern emerging in colonies grown from mixtures of isogenic strains (Fig. S2) [22]. The finding suggests the potential presence of a fitness pressure that selects for matrix producers at the edge of the biofilm at intermediate times, and flagellated cells at later times, removing the need for a single-cell phenotypic response to the local environment to explain the wild-type gene expression pattern.

### Nutrient-dependent fitness landscape explains experimental gene expression pattern

To test whether a stochastic switch coupled to a phenotypic-dependent fitness landscape can explain our observations, we built a reaction-diffusion model where the relative growth rate of flagellated and matrix producing cells depends on the local and instantaneous nutrient concentration. In particular, we hypothesized that flagellated cells grow according to Monod’s law with a constant maximum growth rate, while matrix-producing cells follow Monod’s law with a maximum growth rate that increases linearly with nutrient concentration (Eq. 1). This choice is motivated by the observation that matrix production is energetically demanding and can be interpreted as direct conversion of part of the nutrients into biomass: the more nutrients, the more biomass a single matrix producing cell can generate [23]. Consequently, while some amount of nutrient depletion is necessary to trigger matrix production [20], we expect that matrix producers should grow more slowly than flagellated cells at very low nutrient concentrations and faster, i.e., producing more biomass, in an intermediate range of nutrient concentrations, which overlaps with the initial nutrient availability in our agar plates (Fig. 2, left panel). For wild-type bacteria, a small and constant switching rate between the flagellated and matrix states is also introduced to account for stochasticity in gene expression. The system is then summarized by the following equations:

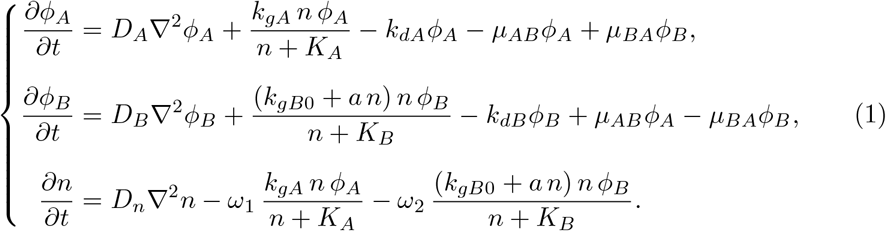

**Fig. 2:**
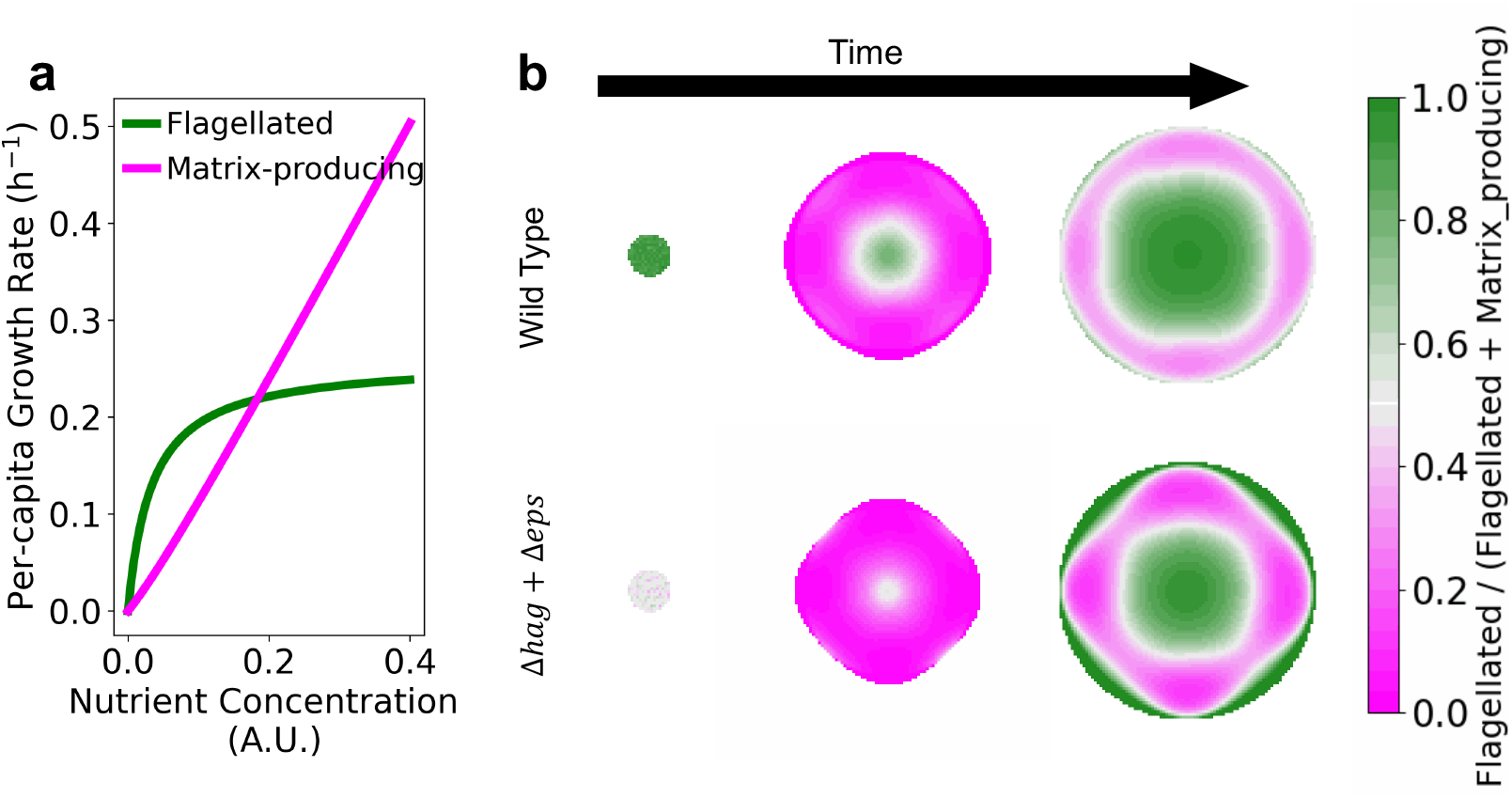
(a) Per-capita growth rate of flagellated (and non-matrix producing) cells (green) and matrix producing (and non-flagellated) cells (magenta) as a function of nutrient concentration in our model (Eqs. 1, Table S3). (b) Resulting gene expression pattern for wild-type (top) at 0 hour, 39 hour, 65 hour and the mixture of knock-out strains (bottom) at 0 hour, 35 hour, 63 hour, respectively.

Here, *ϕ*_*A*_ is the volume fraction of flagellated cells, *ϕ*_*B*_ is the volume fraction of matrix-producing cells, *n* is the nutrient concentration, *D*_*A*_, *D*_*B*_, *D*_*n*_ are their respective diffusion coefficients, *k*_*gA*_ and *k*_*gB*_ = *k*_*gB*0_ + *an* are the maximum growth rates per-capita (*a >* 0), *K*_*A*_, *K*_*B*_ are the half-saturation constants, *k*_*dA*_, *k*_*dB*_ are the death rates, *µ*_*AB*_, *µ*_*BA*_ are the phenotypic switching rates, and *ω*_1_, *ω*_2_ are nutrient-yield coefficients. The switching rates are small constants for wild-type bacteria and set to 0 for knock-out strains. The parameter values are determined by fitting the radial fluorescent intensity profiles of the dual reporter (SI, Fig. S12) and are reported in Table S3, model 1.

Fig. 2 shows that the model is not only able to reproduce the wild-type gene expression pattern on which it is parameterized, but it can also capture the spatio-temporal distribution of the two knock-out strains in the non-switching mixture. In contrast, a constant growth rate coupled to a nutrient-dependent switching rate, which can in principle explain the gene expression pattern of the wild-type, cannot alone reproduce the distribution of knock-out strains (model 2 in Methods, SI and Table S3, Figs. S13 and S14).

Our results suggest that the coordinated gene expression pattern observed in *B. subtilis* biofilms does not require a gene expression response to the local environment at the single-cell level. A simple stochastic phenotypic switch is sufficient to generate enough local diversity to allow the environment to select the appropriate phenotype at the right time and place.

### Phenotypic heterogeneity confers a fitness advantage by shaping and leveraging biofilm’s structure and fluid transport

We then asked whether the presence of a stochastic switch, which enables the wild-type to maintain a phenotypically heterogeneous population, confers some fitness advantage over a non-switching strain in our biofilm conditions. Given the inherent spatial structure of a biofilm grown on solid agar, we started by quantifying fitness as the ability of a population to take over the expanding front from a uniformly distributed inoculum [24, 25], a definition that is routinely used in many biological systems to quantify the evolutionary advantage of populations growing in space [22]. Interestingly, we founf that while the wild-type always out-competes the *hag* knock-out, it is always out-competed by the Δ*eps* mutant (Fig. S3). The result indicates that, from a purely spatial range expansion perspective, lacking matrix production and homogeneously expressing the flagella is a more advantageous strategy than maintaining diversity, since it allows the bacterial biofilm to spread further and faster throughout development.

To test whether phenotypic heterogeneity might provide an advantage under spatial confinement, a situation that is more representative of complex, potentially multispecies biofilm environments [26], we next examined scenarios where one population begins physically enclosed within another (Fig. 3), and we monitored whether the trapped population (*in*) can escape and take over the expanding front of the trapping population (*out*). In most cases, we find the expected result that the *inner* strain is unable to escape and remains physically trapped within the biofilm formed by the *outer* strain (Figs. 3a,c and S4), in agreement with the concept of allele surfing [25] and our model’s predictions (Fig. 3d,f).

**Fig. 3:**
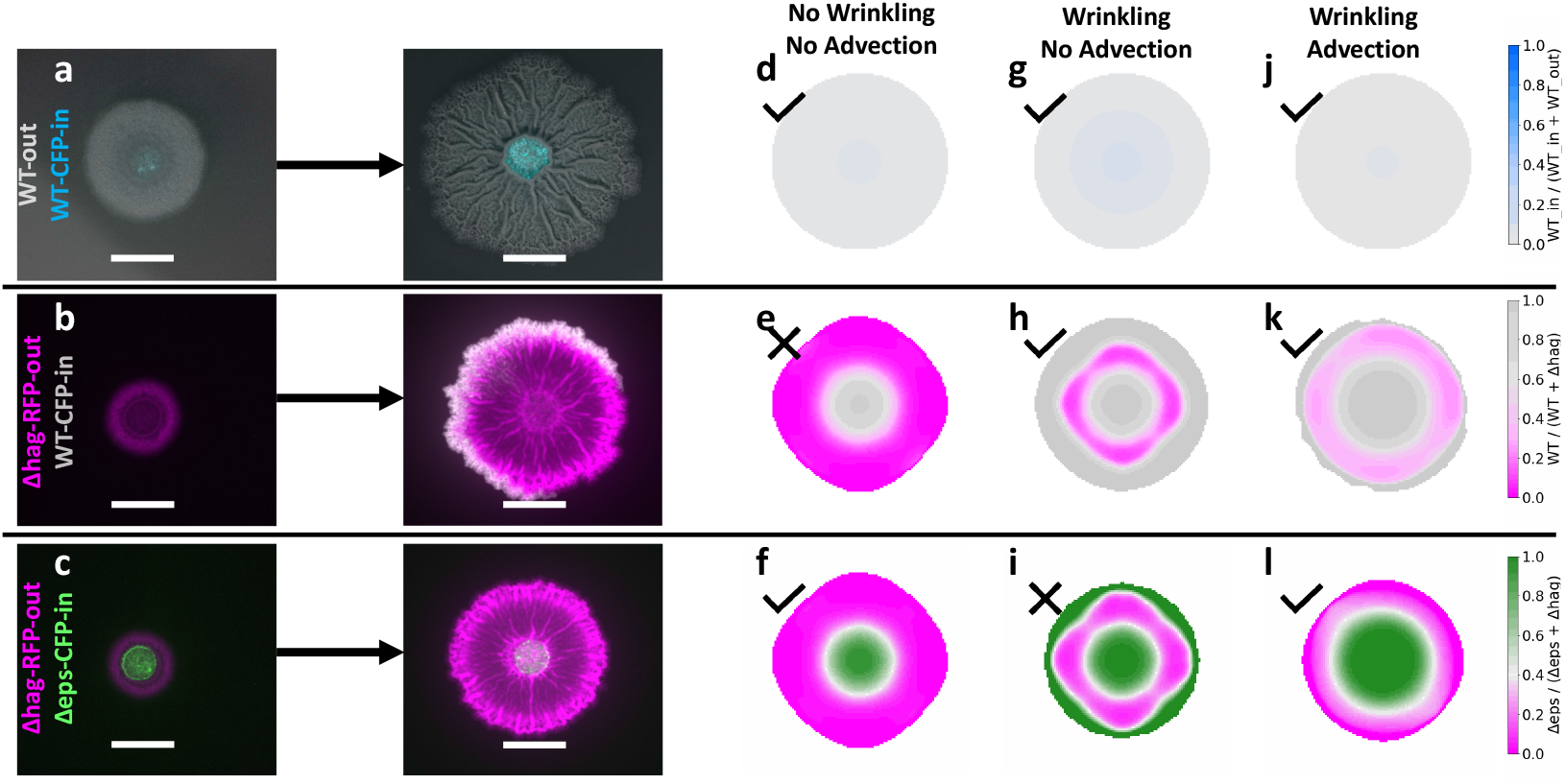
WT cells utilize the wrinkled channels within biofilms produced by non-flagellated *B. subtilis*. Bull’s-eye experiments were set up to assess the escape of inner cells toward the edge of the biofilm. Three competitive setups were tested: WT-out (grey) + WT-CFP-in (cyan) (a), Δ*hag*-RFP-out (magenta) + WT-CFP-in (grey) (b), and Δ*hag*-RFP-out (magenta) + Δ*eps*-CFP-in (green) (c). For each competition, representative merged fluorescent images are shown for day 1 (left) and day 6 (right). Simulation results for analogous competition and initial conditions are shown in (d-l), where (d-f) correspond to model 1 (nutrient-dependent fitness and stochastic switch) (Eqs. 1) at 65 hour, (g-i) correspond to model 3 at 60 hour (increased diffusion of flagellated cells within wrinkle paths), (j-l) correspond to model 4 at 105 hour (advection of flagellated cells within wrinkle paths). (d)(g)(j) for WT-out + WT-in, (e)(h)(k) for Δ*hag*-out + WT-in, and (f)(i)(l) for Δ*hag*-out + Δ*eps*-in.

Unexpectedly, we found one exception to this trend: an initially trapped wild-type population is able to escape a surrounding Δ*hag* biofilm and take over the front (Fig. 3b). A potential explanation for this unexpected result might be the presence of liquid channels that emerge underneath the radial wrinkles of *B. subtilis* biofilms [27]. These channels have been shown to connect the centre to the edge of the colony, enabling nutrient transport across a fully-developed biofilm. We speculated that the same channels could be used by the faster spreading flagellated phenotype of the wild-type as escape routes to overtake the front of the surrounding biofilm. Indeed, we found that when exiting the outer biofilm, wild-type *B. subtilis* strongly expresses the flagellum (Fig. S6) and that an initially *trapped* matrix-only Δ*hag* is unable to escape a surrounding Δ*hag* (Fig. S4a-c). Fluorescence tracking of a wild-type and Δ*hag* cell mixture directly injected into a fully developed biofilm confirmed that both strains can travel the network of channels created by the wrinkles, but wild-type cells do so significantly faster (SI, Fig. S19). The artificial setup of these experiments makes it difficult to generalize any quantification of the relative motility of flagellated vs. non-flagellated cells in the wrinkles to the situation in which cell flow spontaneously occurs during biofilm development. In what follows, therefore, we used this information only to justify our choice to neglect any non-flagellated cell transport phenomenon within the wrinkles and limit the model complexity to the strictly necessary ingredients.

We initially attempted to model cell transport underneath the wrinkles by introducing in our reaction diffusion system radial paths (wrinkles) that allow for increased diffusivity of flagellated bacteria (model 3 in Methods and SI). Once fitted to the dual-reporter gene expression data (Fig. S15), the model successfully predicts the escape of the wild-type strain from the trapping Δ*hag* (Fig. 3h). However, it also predicts that Δ*eps* should be equally able to escape an outer Δ*hag* biofilm (Fig. 3i), in stark contrast to the experimental evidence (Fig. 3c).

To reconcile the opposite result between trapped wild-type and Δ*eps* in our experiments, we built on previous work modelling fluid transport within biofilms [28]: rather than enhancing the diffusivity of flagellated cells in the wrinkles, we introduced an *osmotic pressure* as a function of the local total matrix-producers concentration *ϕ*_*B*_.

Previous studies in *B. subtilis* [28] and *V. cholerae* [10] have provided strong evidence that the likely main driver of osmotic pressure is the ECM, which acts as a sponge and pulls liquid from the agar substrate into the biofilm, causing swelling. Here, we are interested in how this emergent fluid flow couples with cell transport within the channels underneath the wrinkles.

As mentioned above, we neglect the transport of matrix producing cells and assume that only flagellated cells couple to the emergent fluid flow in the channels 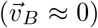. Under quasi-static conditions and Newtonian fluid assumption (model 4 in Methods and SI), this yields a Stokes-type equation for the flagellated cell velocity field

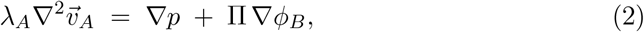

and a frictional coupling between fluid and flagellated cells

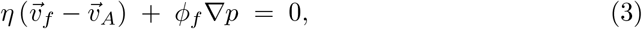

together with incompressibility and volume conservation,

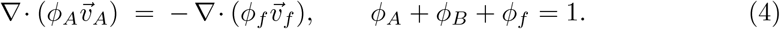

Here, *λ*_*A*_, *η*, and Π control viscosity, cell-fluid friction, and osmotic compressibility, respectively.

Similarly to the previous model with enhanced diffusivity, the transport channels underneath the wrinkles are modelled as radial paths that open after a characteristic time based on experimental observation. Mathematically, we capture this using an indicator *S*(*x, y, t*) that is 1 inside wrinkle regions after opening and 0 otherwise, so that cell transport occurs only where and when channels exist. The resulting model combines growth, death, switching, and diffusion with advection of flagellated cells and fluid:

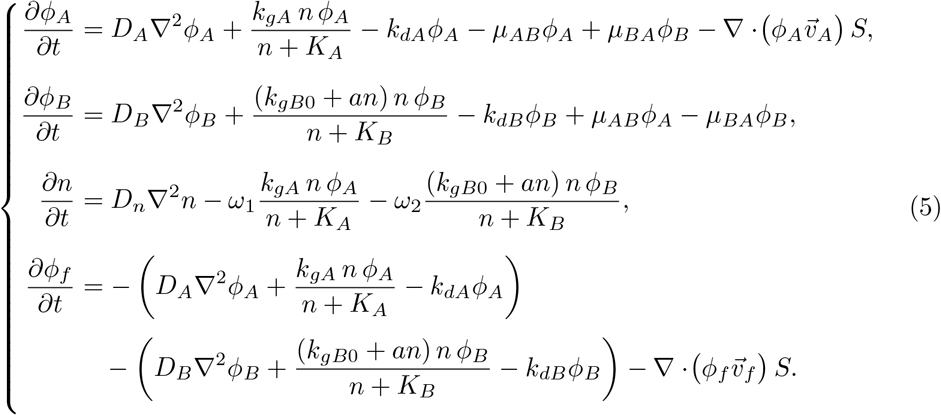

This final model (Fig. S16) is able to explain all the combinations of our *trapping* experiments (Fig. 3j-l and Fig. S4) and highlights how both wrinkle formation and fluid transport are necessary to explain the experimental results. Importantly, the model reveals how phenotypic plasticity is essential to take advantage of the escape routes provided by the wrinkles: even if Δ*eps* mutants can spread faster than the wild-type, only the switching wild-type can successfully escape the surrounding biofilm by maintaining the necessary local diversity to simultaneously generate a pressure gradient (matrix producing cells) and an out-flux of bacteria (flagellated cells). Indeed, trapping a mixture of Δ*hag* and Δ*eps* within a surrounding Δ*hag* biofilm showed no escape of either trapped strain (Fig. S5), underscoring that the phenotypic heterogeneity necessary to observe escape requires to be maintained at small scales and throughout development.

We also point out here that even if the ECM is a shared resource and thus potentially subject to cheating [29], the ECM produced by the outer Δ*hag* does not appear to be exploitable by the trapped Δ*eps*. We think that this is because the osmotic pressure generated by the ECM is transient in nature, as it is proportional to the gradient of the matrix-producing cell density (rather than to the matrix-producing cell density itself), which is significant only at the expansion front (Figs. S16 and S17). In summary, both models and experiments show that a successfully escaping strain must *simultaneously* exhibit both flagella and matrix production, making the phenomenon impervious to potential cheating mechanisms.

### Phenotypic heterogeneity provides the physical ingredients needed for population rescue upon environmental changes

In colonies and biofilms the spatial expansion nature of growth results in the vast majority of spontaneously emerging mutant clones being trapped within the bulk of the colony [30]. This effect is even stronger if the mutations are deleterious, as would be the case for antibiotic resistant clones occurring during biofilm growth in the absence of antibiotic. In a standard smooth colony, these trapped, potentially resistant clones are unable to escape, unless a very high antibiotic concentration is administered that eradicates the surrounding susceptible cells [30].

In a matrix-producing biofilm, the eradication of the bacteria is more challenging, as the ECM provides a physical barrier to antibiotics [31, 32]. While this robust barrier may protect embedded, susceptible cells, it also prevents antibiotic resistant mutants from being unleashed and establishing. Our findings from the previous section suggest that the inherent phenotypic heterogeneity of biofilms and the resulting mechanical and transport properties can equip the bacterial population with escape routes for resistant clones without sacrificing the protection to susceptible cells offered by the ECM.

To investigate the fate of resistant clones within a biofilm undergoing an environmental change, we extended the final model described above (with wrinkles and cell advection) to accommodate a sudden environmental change (e.g., antibiotic administration) that provides a significant growth advantage to the trapped strain (resistant clone), while limiting the growth of the surrounding strain (susceptible background) without altering its structure (SI). Our model predicts that an initially trapped resistant clone would be able to escape the surrounding biofilm by flowing underneath the wrinkles (Fig. 4f). In contrast, if the trapped clone lacks either matrix production (Δ*eps*, Fig. 4g) or flagella (Δ*hag*, Fig. 4h), escape cannot occur, because cell transport is prevented either by lack of osmotic pressure (Δ*eps*) or by lack of flagellated cells (Δ*hag*).

**Fig. 4:**
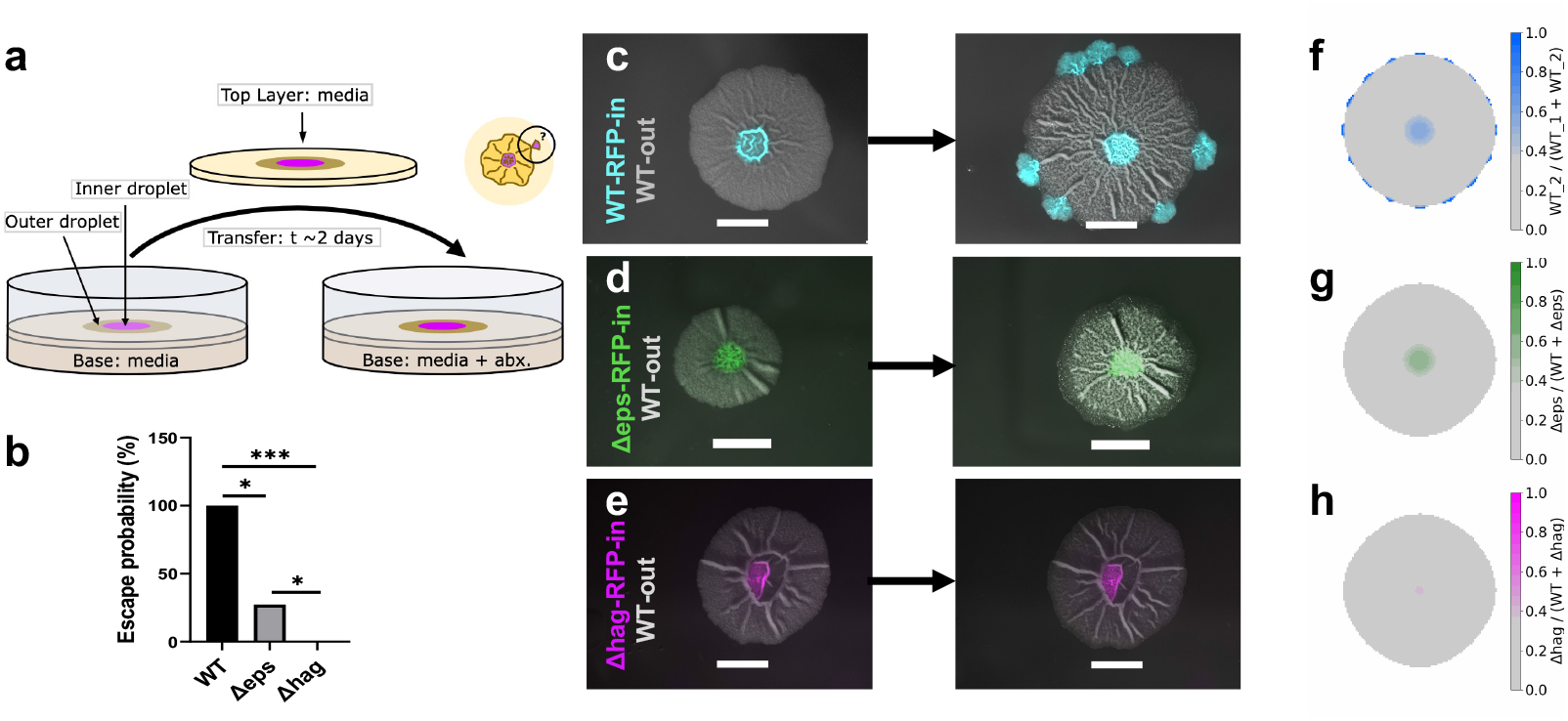
Initially trapped WT cells emerge at the edge of an expanding biofilm in association with wrinkles. (a) Schematic of the experimental setup. (b) Column graph showing escape probabilities of WT, Δ*eps*, and Δ*hag* strains in the presence of wrinkles within a WT biofilm. Statistical significance was assessed using Fisher’s exact test with pairwise comparisons in GraphPad. ns = P *>* 0.05; * = P ≤ 0.05; ** = P ≤ 0.01; *** = P ≤ 0.001; **** = P ≤ 0.0001. (c) When the outer droplet was a non-fluorescent WT biofilm-forming strain and the inner droplet was an analogous strain (WT-RFP-PhleoR) expressing RFP and resistant to phleomycin, fluorescent cell domains appeared at the biofilm edge and co-localized with the termini of radial wrinkles. (d) When the inner droplet contained Δ*eps*-RFP-PhleoR cells (flagellated but unable to produce EPS matrix), the probability of escape was reduced, and most replicates showed no escape events. (e) A flagellum-null strain (Δ*hag*-RFP-PhleoR) was unable to escape, even in the presence of wrinkles, due to its inability to swim through the wrinkled channels. Scale bars: 5000 *µ* m. Corresponding simulation results of our wrinkling-advection model at 25 hours after transfer process are shown in (f-h), where (f) for WT-out-WT-in, (g) for WT-out-Δ*eps*-in, and (h) for WT-out-Δ*hag*-in.

To verify the model’s predictions, we devised a new protocol that allows changing the chemical environment of the biofilm without altering its spatial structure (Fig. 4a, SI). In brief, we inoculated two strains so that an inner small droplet of a resistant clone (inner droplet) was encompassed within a larger droplet of a resident susceptible population (outer droplet) on a thin agar slab containing MSgg media. At the time of inoculation, the slab was placed on top of a thicker layer of agar containing the same media. After two days during which wrinkles emerged, the thinner slab was transferred to a new base layer containing antibiotic, inducing a sudden environmental change without manipulation of the spatial structure of the biofilm.

In agreement with the model, we found that a resistant wild-type strain embedded in a susceptible wild-type strain can consistently escape the surrounding population and establish at the front (Fig. 4b,c). The locations where the resistant clone escapes were reproducibly aligned with the end-points of the radial wrinkles of the surrounding biofilm (Fig. S8), supporting our hypothesis that the channels underneath the wrinkles are used by the trapped cells to traverse the biofilm and reach the front. Indeed, we found that when wrinkles were not present in the surrounding biofilm, either because the surrounding strain does not produce the necessary matrix, or because the medium was treated to prevent proper wrinkle formation, escape did not occur (Fig. S9a,e). Similarly, if biofilm transfer occurs too early (∼1 day), before the wrinkles are properly formed, escape does not occur (Fig. S9b).

Also in agreement with the model’s predictions, we found that neither knockout mutant strains could successfully escape (Fig. 4d,e). In line with the model’s assumptions, this result confirms that the cell transport process we observed cannot be explained by growth or chemotaxis alone, since both phenomena would predict a successful escape of the Δ*eps* strain. In further support of this statement, we also found that wild-type cells can successfully escape even when the biofilm is transferred to an antibiotic pad without media supplementation (Fig. S9d), so where additional growth or chemotactic cues for the trapped strain are minimal.

## Discussion

Coordinated gene expression patterns in biofilms have been documented across different bacterial species and conditions. In *B. subtilis*, in particular, several studies have carefully investigated the sequence of events that leads to sporulation, providing unprecedented data on the regulation, stochastic fluctuations, and dynamics of the gene circuit that determines the matrix producing vs. flagellated state of the cell. The resulting picture is partially controversial. On one side, several studies, typically population-based, show evidence of an environmentally controlled switch, where cell fate is determined by the presence of certain environmental cues [17, 20, 33]. On the other side, several single cell analyses have demonstrated that cell states change stochastically even under perfectly homogeneous conditions [13, 18]. These results raise two intriguing questions: (i) is an environment-responsive switch necessary to explain the gene expression patterns observed in biofilms? and (ii) does maintaining a stochastic switch between flagellated and matrix producing cells confer a fitness advantage to the population?

When monitoring *B. subtilis* biofilms grown on solid-air interfaces, we found that a biofilm containing an initial mixed population of knock-out strains that are unable to switch between functional phenotypes spontaneously forms a spatio-temporal distribution that qualitatively matches the gene expression patterns of wild-type biofilms (Fig. 1). More specifically, we observed that cells that cannot produce matrix are depleted at the edge early in biofilm development, while cells that cannot form the flagellum are depleted at the edge later in development. This result suggests the presence of a fitness landscape that confers a growth rate (dis)advantage to matrix producing vs. flagellated cells at different points in space and time. Given the geometry of the pattern, we hypothesized that such growth rate (dis)advantage is coupled with nutrient concentration. We found that a reaction-diffusion model with such fitness landscape and a purely stochastic matrix-flagellated switch can explain the gene expression pattern of wild-type biofilms and the strain spatio-temporal distribution in the mixed biofilm without the need of an environment-responsive switch (Fig. 2).

This finding puts forward the hypothesis that some gene expression patterns observed in biofilms might be simply explained by the environment selecting for specific phenotypes at given points in time and space, without requiring individual cells to actively sense and respond to the environment by activating certain gene pathways. An analogous observation was made when monitoring the proliferation of flagellated versus matrix producing cells subjected to electrical stimuli in microfluidic devices [34]. Discriminating whether an observed gene expression pattern in the biofilm is the result of individual cells responding to the environment or a different growth rate between populations of cells expressing different genes is challenging at the population level. Single-cell data that simultaneously monitor growth rate and switching rate in a range of environmental conditions is key to differentiate between these scenarios, and we do not exclude the possibility that both rates might be influenced by the environment. More data in this direction will help answer fundamental questions on the evolution of phenotypic heterogeneity and phenotypic switches, e.g., whether stochastic switches are evolutionary easier to achieve than responsive switches or whether a stochastic switch confers unforeseen advantages compared to a responsive switch.

The observation that the phenotype selected by the environment at the edge of a fully developed biofilm is flagellum production begs the question of why selection for matrix production at the outer edge occurs at intermediate times. Indeed, we found that the Δ*eps* knock-out spreads further and faster than the wild-type, suggesting that a homogeneous non-matrix-producing population would be more competitive than the stochastically switching wild-type in terms of colonizing virgin territory.

We found a potential answer to this question when we changed our measure of competitive advantage from pure growth to survival to environmental changes. It is well-known that in biofilms, as in any other spatially expanding population, the vast majority of spontaneous mutant clones is spatially trapped within the bulk of the population, providing a reservoir of genetic variation that cannot be unleashed unless the surrounding cells are physically removed. In biofilms, specifically, such eradication is even more challenging due to the presence of the ECM. How can the biofilm then take advantage of mutant clones that can be beneficial in new environments while maintaining the protection provided by the ECM? Combining experiments and physical models, we found that a solution to this problems comes from an unexpected and elegant interplay between the phenotypic heterogeneity and the biofilm’s physical properties. Matrix production, which our environment favours early in development, is essential not only to provide a physical barrier for the enclosed cells but also to generate the mechanical instabilities that lead to wrinkle formation, which, in turn, result in a network of channels akin to a vascular system. This architecture, coupled with additional matrix production, appears to be used by the biofilm to adsorb liquid from the substrate and push flagellated cells that are initially deep inside the biofilm outwards to the edge. Spontaneously trapped clones can then have a route to disperse without any disruption to the ECM and continue the expansion in a new environment that prevented the growth of the original biofilm (Fig. 4). In summary, early matrix production is necessary for the formation and pumping of the *biofilm vascular system*, and flagellum expression is necessary for utilization of this system and eventual population rescue. A stochastic switch between flagellated and matrix-producing states ensures that subpopulations of both phenotypes are always available to take on one function or the other depending on the local physical environment that they themselves generate or encounter [35, 36], so as to maximize biofilm’s resilience in these conditions, protect the population from potential cheating mutants and external invaders, and generally enhance adaptability.

Our model provides a potential explanation for the physical mechanism underlying the required cooperation between matrix producing and flagellated cells to observe the escape of trapped clones; however, alternative mechanisms can also be envisioned. The outflux of cells coupled with the influx of fluid might be caused by the propensity of flagellated cells to swim *upstream* when in proximity to an interface [37] or by surface tension interactions between the ECM and the surfactant molecules produced by flagellated cells [38]. In addition, matrix production might promote fluid flow by *rebuilding* collapsed channels, rather than through osmotic pressure. Controlled experiments, possibly in microfluidic devices, would be instrumental to pinpoint the exact mechanism behind the interplay between matrix production and motility that our experiments reveal, and explore its applicability to biofilms growing in different environments and conditions.

## Methods

### Experimental methods

#### Bacterial strains and media

The strains used are biofilm-forming *B. subtilis* strains derived from NCIB 3610. WT *B. subtilis* (3610 and 3A38) and their knock-out mutants (Δ*hag* and Δ*eps*) are listed in Table S1. Details about the genetic engineering methods employed are reported in the SI file (Table S2).

Strains were grown in LB (Miller’s lysogeny broth base, Invitrogen) or the defined MSgg medium [39], which is known to promote biofilm growth (5 mM phosphate buffer pH 7.0; 100 mM MOPS buffer pH 7.0, adjusted with NaOH; 0.5% (v/v) glycerol; 0.5% (w/v) monosodium glutamate; 2 mM MgCl_2_; 0.7 mM CaCl_2_; 50 *µ*M MnCl_2_; 50 *µ*M FeCl_3_; 1 *µ*M ZnCl_2_; 2 *µ*M thiamine HCl; 50 *µ*g/mL tryptophan; 50 *µ*g/mL phenylalanine).

#### Mixed droplets

For mixed droplet experiments, overnight-grown cultures were washed three times with phosphate buffer saline. Cell suspensions were adjusted to an OD_600_ of 1.0 and mixed at a 1:1 ratio to prepare a competitive strain mixture [40]. MSgg agar plates were prepared one day prior to the experiment by autoclaving 1.5% (w/v) agar (VWR Chemicals) with MOPS buffer, phosphate buffer, and glycerol. After cooling, the remaining components were added from sterile-filtered stock solutions. Fresh monosodium glutamate stock was prepared on the day of use. A total of 25 mL of molten MSgg agar was poured into standard 90-mm Petri plates. Once solidified, the plates were dried overnight at room temperature. The following day, the plates were further dried for 1 h in a laminar flow hood. Finally, 1 *µ*L of the cell suspension (individual or mixed droplet) was inoculated onto the MSgg agar plates, which were then sealed with masking tape and incubated at 30 °C.

#### Bull’s eye experiments - no transfer

For the bull’s-eye experiments, MSgg plates were prepared in the same way. Overnight cultures were adjusted to an OD_600_ ≈ 1, as described earlier. For the outer droplets, 2 *µ*L of the resident strain was pipetted onto the surface of the MSgg agar and allowed to dry for ∼ 3 minutes. Once the outer droplet had dried, a 0.5 *µ*L inoculation of the test strain was placed in the centre of the outer droplet, completely surrounded by it, and allowed to dry for *∼* 1 minute. Plates were sealed and incubated at 30 °C.

#### Bull’s eye experiments - with transfer

For the bull’s-eye transfer experiments, MSgg agar was poured into 128 × 86 mm rectangular plates (Nunc OmniTray Single-Well Plate, Thermo Scientific), and 10 mL was poured into 90-mm Petri plates. After drying, a square portion of the thinner MSgg agar layer from the 90-mm Petri plates was aseptically transferred to each of the thicker MSgg agar plates. Strains were inoculated as in the bull’s-eye experiment. After two days, the biofilm was imaged, and the top layer of MSgg agar was transferred to another base layer of MSgg agar containing antibiotic selection (10 *µ*g/mL phleomycin, Sigma-Aldrich), which provided a growth advantage to the inner strain (fig. S6). This plate had been prepared the previous day by drying overnight on the bench and then in the laminar hood, as described above. Biofilm development was monitored by time-lapse imaging (every 40 minutes) or by regular still images using a fluorescence microscope with a 0.5× objective as described below. Statistical analysis for the transfer experiment (Fig. 4b) was performed using GraphPad Prism 9. P-values are presented following GraphPad’s significance notation: 0.1234 (ns), 0.0332 (*), 0.0021 (**), 0.0002 (***), ¡0.0001 (****).

#### Biofilm imaging

Biofilms were imaged using an upright Zeiss Axio Zoom.V16 stereo microscope equipped with a Zeiss PlanApo Z 0.5 ×/0.125 FWD 114-mm objective and a Plan-NeoFluar Z 2.3 ×/0.57 FWD 10.6-mm objective. The former was used for full biofilm imaging, while the latter was used to image the concentrated mixture of cells within the biofilm channels or for single-cell imaging. During imaging, samples were incubated at 30 °C in a custom-made incubation chamber enclosing the entire microscope setup. For time-lapses, images were taken every 40 minutes from above through a thin transparent glass heated to 32 °C to prevent condensation. For time-lapses taken after the introduction of the concentrated mixture of cells to the biofilm (see SI), images were taken every 24 seconds, and imaged from below the biofilm through the agar. The brightness and contrast in the images were adjusted using the FIJI distribution (Version 1.54m) of ImageJ [41].

### Modelling

We developed four reaction-diffusion(–advection) models to identify the minimal ingredients necessary to qualitatively explain the experimental results. We denote *ϕ*_*A*_(*x, y, t*) and *ϕ*_*B*_(*x, y, t*) as the local volume fractions of flagellated and matrix-producing cells, respectively, and *n*(*x, y, t*) as the local nutrient concentration. Local volume fraction of fluid *ϕ*_*f*_ is added to model 4 to better explain the advection term. Unless stated otherwise, we assume: (i) Monod growth; (ii) constant death rates *k*_*dA*_, *k*_*dB*_; (iii) stochastic switching with rates *µ*_*AB*_ (from flagellated to matrix), and *µ*_*BA*_ (from matrix to flagellated); (iv) constant initial nutrient concentration; (v) zero-flux boundary conditions.

The different variants are briefly summarized here, with further details, parameters fitting procedure, and parameter values reported in the SI.

#### Model 1: stochastic switch and nutrient-dependent fitness

We assume a small stochastic switch between flagellated and matrix state and a maximum growth rate that is constant for flagellated cells and nutrient-dependent for matrix cells. The mathematical equations describing the model are reported in eq. 1 and more details can be found in the SI.

#### Model 2: nutrient-dependent switch and constant fitness

Here, we model a switch rate that depends on the local environment, specifically nutrient concentration, favoring the switch from flagellated to matrix state at high nutrient concentration, and from matrix to flagellated state at low nutrient concentration. Maxima growth rates are held constant. This leads to the following system of PDEs:

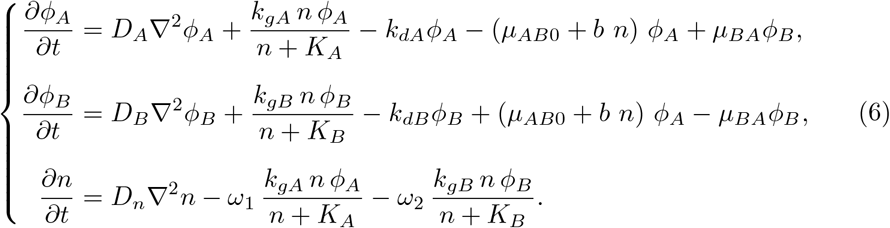

#### Model 3: stochastic switch, nutrient-dependent fitness, and enhanced diffusivity of flagellated cells within wrinkles

This model accommodates the liquid channel emerging underneath the biofilm wrinkles via the modification of the diffusion term for flagellated cells in model 1, eq. 1. Within the wrinkles, defined as radial paths from the edge of initial droplet (the inner circle of biofilm) to the edge of biofilm, 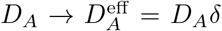, where *δ* = 1 before the wrinkles open (*t < t*_0_), and increases to a value larger than 1 for *t* ≥ *t*_0_ (parameters reported in Table S3).

#### Model 4: stochastic switch, nutrient-dependent fitness, and flagellated cells advection within wrinkles

In this model, we accommodate cell transport within the wrinkles by introducing an advection term for the flagellated cells that is coupled to a fluid flux driven by matrix production. The mathematical formulation is reported in eq. 5. Importantly, the model assumes that the osmotic pressure generate by the EPS is proportional to the density of matrix producing cells *ϕ*_*B*_ and that matrix cells, by sticking to the walls of the wrinkles, are unaffected by the fluid flux. Details of the derivation are reported in the SI.

#### Model fitting and parameter estimation

All models are parameterized by fitting the radial profiles of YFP and RFP fluorescence intensity from the dual-reporter time-lapse to the spatio-temporal profiles for *ϕ*_*A*_ and *ϕ*_*B*_ predicted by the models (see also SI). To convert fluorescence intensity (output of the experiments) to cell density and cell volume fraction (output of the model), we perform single-cell imaging analysis on cells sampled from a mature dual-reported biofilm, as reported in the SI.

## Data and code availability

All the Jupyter notebook codes used to generate the modelling figures in this work can be found here: https://github.com/FuscoLab/2025 motile matrix plasticity.

## Acknowledgments

This work was supported by a Research Grant from the Human Frontier Science Program: doi.org/10.52044/HFSP.RGY00572022.pc.gr.153591. AS and DF also acknowledge support from an ERC Starting Grant/UKRI Horizon Europe Guarantee (EP/Y030141/1). We thank Munehiro Asally, Joseph Larkin, James Locke and Agnese Seminara for kindly providing some of the strains used in this work.

## Author contributions

AM, NK, and DF conceptualized the project and designed the experiments. AM engineered the strains. AM, NK and JK carried out the experiments. YD and DF developed the model. YD implemented the model and analyzed the gene expression profiles. AKV and LRP analyzed the cell transport data. AS analyzed the single cell microscopy data. CT, LRP and DF acquired funding. DF led the manuscript writing, and all authors contributed and approved the final version.

## Supplementary information

Supplementary Information is available for this paper.

## References

[1] Obando, M. C. & Serra, D. O. Dissecting cell heterogeneities in bacterial biofilms and their implications for antibiotic tolerance. Current opinion in microbiology 78, 102450 (2024).

[2] Serra, D. O. & Hengge, R. A c-di-gmp-based switch controls local heterogeneity of extracellular matrix synthesis which is crucial for integrity and morphogenesis of escherichia coli macrocolony biofilms. Journal of molecular biology 431, 4775–4793 (2019).

[3] Nadezhdin, E., Murphy, N., Dalchau, N., Phillips, A. & Locke, J. C. Stochastic pulsing of gene expression enables the generation of spatial patterns in bacillus subtilis biofilms. Nature communications 11, 950 (2020).

[4] Gestel, J. V., Vlamakis, H. & Kolter, R. Division of labor in biofilms: the ecology of cell differentiation. Microbial Biofilms 67–97 (2015).

[5] Yannarell, S. M. et al. Extensive cellular multi-tasking within bacillus subtilis biofilms. MSystems 8, e00891–22 (2023).

[6] Grimbergen, A. J., Siebring, J., Solopova, A. & Kuipers, O. P. Microbial bethedging: the power of being different. Current opinion in microbiology 25, 67–72 (2015).

[7] Kampf, J. et al. Selective pressure for biofilm formation in bacillus subtilis: differential effect of mutations in the master regulator sinr on bistability. MBio 9, 10–1128 (2018).

[8] Lowery, N. V., McNally, L., Ratcliff, W. C. & Brown, S. P. Division of labor, bet hedging, and the evolution of mixed biofilm investment strategies. MBio 8, 10–1128 (2017).

[9] Smith, P. & Schuster, M. Public goods and cheating in microbes. Current biology 29, R442–R447 (2019).

[10] Yan, J., Nadell, C. D., Stone, H. A., Wingreen, N. S. & Bassler, B. L. Extracellular-matrix-mediated osmotic pressure drives vibrio cholerae biofilm expansion and cheater exclusion. Nature communications 8, 327 (2017).

[11] Elowitz, M. B., Levine, A. J., Siggia, E. D. & Swain, P. S. Stochastic gene expression in a single cell. Science 297, 1183–1186 (2002).

[12] Rossi, E., Paroni, M. & Landini, P. Biofilm and motility in response to environmental and host-related signals in gram negative opportunistic pathogens. Journal of Applied Microbiology 125, 1587–1602 (2018).

[13] Lord, N. D. et al. Stochastic antagonism between two proteins governs a bacterial cell fate switch. Science 366, 116–120 (2019).

[14] Vlamakis, H., Aguilar, C., Losick, R. & Kolter, R. Control of cell fate by the formation of an architecturally complex bacterial community. Genes & development 22, 945–953 (2008).

[15] Branda, S. S., González-Pastor, J. E., Ben-Yehuda, S., Losick, R. & Kolter, R. Fruiting body formation by bacillus subtilis. Proceedings of the National Academy of Sciences 98, 11621–11626 (2001).

[16] Krishnan, N., Knight, J., Tropini, C., Ruiz Pestana, L. & Fusco, D. The what, when, where, and why of wrinkly morphology in biofilms. Biophysics reviews 6 (2025).

[17] Srinivasan, S. et al. Matrix production and sporulation in bacillus subtilis biofilms localize to propagating wave fronts. Biophysical Journal 114, 1490–1498 (2018).

[18] Norman, T. M., Lord, N. D., Paulsson, J. & Losick, R. Memory and modularity in cell-fate decision making. Nature 503, 481–486 (2013).

[19] Dannenberg, S., Penning, J., Simm, A. & Klumpp, S. The motility-matrix production switch in bacillus subtilis—a modeling perspective. Journal of Bacteriology 206, e00047–23 (2024).

[20] Zhang, W. et al. Nutrient depletion in bacillus subtilis biofilms triggers matrix production. New Journal of Physics 16, 015028 (2014).

[21] Asally, M. et al. Localized cell death focuses mechanical forces during 3d patterning in a biofilm. Proceedings of the National Academy of Sciences 109, 18891–18896 (2012).

[22] Hallatschek, O. & Nelson, D. R. Life at the front of an expanding population. Evolution 64, 193–206 (2010).

[23] Xavier, J. B. & Foster, K. R. Cooperation and conflict in microbial biofilms. Proceedings of the National Academy of Sciences 104, 876–881 (2007).

[24] Korolev, K. S. et al. Selective sweeps in growing microbial colonies. Physical biology 9, 026008 (2012).

[25] Gralka, M. et al. Allele surfing promotes microbial adaptation from standing variation. Ecology letters 19, 889–898 (2016).

[26] Porter, R. et al. On the growth and form of bacterial colonies. Nature Reviews Physics 1–19 (2025).

[27] Wilking, J. N. et al. Liquid transport facilitated by channels in bacillus subtilis biofilms. Proceedings of the National Academy of Sciences 110, 848–852 (2013).

[28] Seminara, A. et al. Osmotic spreading of bacillus subtilis biofilms driven by an extracellular matrix. Proceedings of the National Academy of Sciences 109, 1116–1121 (2012).

[29] Martin, M. et al. Cheaters shape the evolution of phenotypic heterogeneity in bacillus subtilis biofilms. The ISME journal 14, 2302–2312 (2020).

[30] Fusco, D., Gralka, M., Kayser, J., Anderson, A. & Hallatschek, O. Excess of mutational jackpot events in expanding populations revealed by spatial luria– delbrück experiments. Nature communications 7, 12760 (2016).

[31] Tseng, B. S. et al. The extracellular matrix protects p seudomonas aeruginosa biofilms by limiting the penetration of tobramycin. Environmental microbiology 15, 2865–2878 (2013).

[32] Shree, P., Singh, C. K., Sodhi, K. K., Surya, J. N. & Singh, D. K. Biofilms: Understanding the structure and contribution towards bacterial resistance in antibiotics. Medicine in Microecology 16, 100084 (2023).

[33] Chou, K.-T. et al. A segmentation clock patterns cellular differentiation in a bacterial biofilm. Cell 185, 145–157 (2022).

[34] Comerci, C. J. et al. Localized electrical stimulation triggers cell-type-specific proliferation in biofilms. Cell Systems 13, 488–498 (2022).

[35] Kearns, D. B. & Losick, R. Cell population heterogeneity during growth of bacillus subtilis. Genes & development 19, 3083–3094 (2005).

[36] Steinberg, N. et al. The extracellular matrix protein tasa is a developmental cue that maintains a motile subpopulation within bacillus subtilis biofilms. Science signaling 13, eaaw8905 (2020).

[37] Kaya, T. & Koser, H. Direct upstream motility in escherichia coli. Biophysical journal 102, 1514–1523 (2012).

[38] Li, Y. et al. Self-organized canals enable long-range directed material transport in bacterial communities. Elife 11, e79780 (2022).

[39] Branda, S. S., Chu, F., Kearns, D. B., Losick, R. & Kolter, R. A major protein component of the bacillus subtilis biofilm matrix. Molecular microbiology 59, 1229–1238 (2006).

[40] Dragoš, A. et al. Division of labor during biofilm matrix production. Current Biology 28, 1903–1913.e5 (2018). URL https://www.sciencedirect.com/science/article/pii/S0960982218305189.

[41] Schindelin, J. et al. Fiji: an open-source platform for biological-image analysis. Nature methods 9, 676–682 (2012).

