## Supplementary Information for "Phenotypic plasticity shapes biofilm’s structure and fluid transport enhancing resilience"

#### 1 Strain engineering

Flagellar knock-out mutants on WT-CFP and WT-RFP background were constructed using the pTS157 plasmid (courtesy of James Locke lab), which replaces *hag* with the kanamycin resistance cassette. Genomic DNA of *B. subtilis* 3610 *epsA-O::tet* was used for knocking out the *epsA-O* operon from the WT-CFP, WT-RFP and WT-RFP-PhleoR.

A standard one-step transformation procedure was used to construct the *epsA-O* knock-out mutants [1]. A two-step transformation protocol was followed for constructing the *hag* knock-outs. For the two-step protocol, in brief, SPC medium (50 mL) was prepared with sterile 5 mL 10× SMS, 1 mL 25% glucose, 2 mL 5% yeast extract, 1.25 mL 1% casamino acids, 0.8 mL amino acid mix (2.5 mg/mL; tryptophan required), and water to 50 mL. SPII medium (50 mL) contained 5 mL 10× SMS, 1 mL 25% glucose, 1 mL 5% yeast extract, 0.5 mL 1% casamino acids, 0.8 mL amino acid mix, 0.5 mL 50 mM $CaCl_2$, 0.25 mL 1 M $MgSO_4$, and water to 50 mL. SPC and SPII were filter sterilized and prepared fresh. ME solution (2 mL) consisted of 1.76 mL water, 0.2 mL 10× SMS, and 0.04 mL 100 mM EGTA (pH 8.0, adjusted until dissolved), and was sterilized by filtration immediately before use. For 10× SMS (250 mL), 5 g $(NH_4)_2SO_4$, 35 g $K_2HPO_4$, 15 g $KH_2PO_4$, 2.5 g $Na_3$-citrate·$2H_2O$, and 0.5 g $MgSO_4$·$7H_2O$ were dissolved and filter sterilized. LB agar (1 L) was prepared with 10 g tryptone, 5 g yeast extract, 10 g NaCl, 1 mL 1 M NaOH, and 15 g agar, followed by autoclaving.

Overnight cultures (≤16–18 h) were grown in LB. One milliliter of culture was pelleted at 5000 rpm for 5 min and resuspended in 5 mL SPC medium in a 50 mL tube. Cells were incubated at 37 °C until reaching constant OD (typically $\sim$ 6 h). Cultures were then diluted 1:10 into pre-warmed SPII (1 mL cells + 9 mL SPII) and incubated for 90 min at 37 °C while shaking. Cells were pelleted at 4000 rpm for 3 min and resuspended in 1 mL supernatant. 100 $\mu$L of competent culture was mixed with 100 $\mu$L ME solution and 1–2 $\mu$g DNA (≤10 $\mu$L volume). Mixtures were incubated at 37 °C while shaking for 30–60 min before plating on selective agar. Construction of positive mutants was verified using PCR screening with corresponding primers (Table S2). Bright-field images of mature biofilms of these strains grown on MSgg agar plates are shown in fig. S1.

**Table S1** Strains and plasmids used in this study. All strains are either in the wild-type *B. subtilis* 3610 or 3A38 background.

| Strain/ Plasmid | Genetic information | Source |
|---|---|---|
| WT | *B. subtilis* NCIB 3610 | |
| WT 3a38 | *B. subtilis* BGSC 3A38 | |
| Dual-reporter | *B. subtilis* NCIB 3610 sacA::Phag-yfp, amyE::PtasA-tsr-mcherry | Munehiro Asally |
| WT-CFP | *B. subtilis* 3A38 amyE::PvegS-mTurquoise2 | Joseph Larkin |
| WT-RFP | *B. subtilis* 3A38 amyE::PvegS-mScarlet | Joseph Larkin |
| Δ*hag* | *B. subtilis* NCIB 3610, hag::tet | Agnese Seminara |
| Δ*hag*-CFP | *B. subtilis* 3A38 amyE::PvegS-mTurquoise2, hag::Km | this paper |
| Δ*hag*-RFP | *B. subtilis* 3A38 amyE::PvegS-mScarlet, hag::Km | this paper |
| Δ*eps*-CFP | *B. subtilis* 3A38 amyE::PvegS-mTurquoise2, epsA-O::Tet | this paper |
| Δ*eps*-RFP | *B. subtilis* 3A38 amyE::PvegS-mScarlet, epsA-O::Tet | this paper |
| Δ*eps* | *B. subtilis* 3610 (HV1182); epsA-O::tet | Agnese Seminara |
| WT-RFP-PhleoR | *B. subtilis* 3A38 ppsb::PtrpE-mCherry-PhleoR | James Locke/Teresa Saez |
| Δ*hag*-RFP-PhleoR | *B. subtilis* 3A38 ppsb::PtrpE-mCherry-PhleoR, hag::Km | James Locke/Teresa Saez |
| Δ*eps*-RFP-PhleoR | *B. subtilis* 3A38 ppsb::PtrpE-mCherry-PhleoR, epsA-O::tet | this paper |
| pTS157 | Plasmid pTS157 having 5'-hag-Km-hag-3' cassette | James Locke/Teresa Saez |

**Table S2** Primers and PCR conditions used in this study. NEB Q5 enzyme was used for PCR reactions. The $\Delta epsA-O$ mutant verification primer set should bind upstream of epsA-O operon and depsVerRn should bind with TetR gene. The $\Delta hag$ mutant verification primer set should bind hag gene and delhagKmR should bind KmR gene. AT stands for annealing temperature, ET for elongation time and NC for number of cycles.

| Used in verification | Primer | Sequence (5'-3') | Product size | AT | ET | NC |
|---|---|---|---|---|---|---|
| $\Delta epsA-O$ mutants | depsVerF<br>depsVerR | CAAGCAATCCTCGGACTGGC<br>CCAGTTTGTACTCGCAGGTGG | ∼ 1400 bp | 69 °C | 1 min | 30 |
| $\Delta hag$ mutants | delhagKmF<br>delhagKmR | ATGGCAGAGCCATTTGAAAAGTC<br>ATGGAGTGTCTTCTTCCCAGT | ∼ 1250 bp | 66 °C | 1 min | 30 |

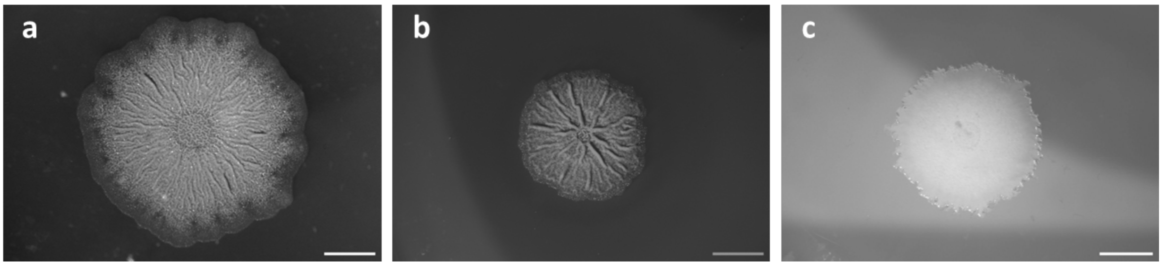


**Fig. S1** Brightfield images of mature biofilms formed by *B. subtilis* 3610 strains: (a) wild type (WT) (day 6), (b) non-flagellated $\Delta hag$-CFP (day 7), and (c) non-flagellated $\Delta eps$-CFP (day 7). Scale bars represent 5000 $\mu m$.

### 2 Single-cell imaging

To convert fluorescence intensity of the dual-reporter strain to number of cells expressing either matrix or motility genes, we performed the following single-cell imaging protocol. The dual reporter strain was grown for one day on an MSgg agar plate to allow biofilm development. A loopful of biofilm was scraped and resuspended in 100 $\mu$l PBS buffer. The biofilm sample was vigorously vortexed to obtain a homogeneous cell suspension. Small agarose (1.5%) pads were prepared using a silicone mold on a glass coverslip. One microliter of the cell suspension was placed on the agarose pad and imaged under the microscope. Several snapshots were taken of both the single cells or the original biofilm, and these were analyzed as described in the following section.

#### 2.1 Single-cell image analysis

##### 2.1.1 Segmentation

The fluorescence images were segmented using Cellpose 2 [2], which creates one labeled 'mask' for each cell in the image (approximately 26,000 cells in total across all images). We then calculate:

- The size of the cell, equal to the area of the mask (fig. S10a,d).
- The total fluorescent expression of the cell, equal to the sum of the values of all pixels inside the mask, in the YFP or RFP channel respectively (fig. S10b,e).
- The fluorescent concentration of the cell, equal to the total fluoresecent expression divided by the area (fig. S10c,f).

In some cases, usually where cells were clustered together in an otherwise sparse environment, Cellpose assigned a single label to multiple cells. To correct for this, the median area of all YFP expressing cells (and separately RFP expressing cells) was calculated (dashed black line in fig. S10a,d), and any object more than twice this area was discarded from further analysis (black bars in fig. S10a-f). The remaining data were then averaged to calculate a mean area, mean total fluorescence, and mean fluorescence concentration for YFP and separately RFP-expressing cells.

##### 2.1.2 Fluorescence ratio $\alpha$ and intensity-volume ratio $\beta$

To inform the model of the value of parameter $\alpha$ (intensity ratio of a cell expressing matrix over a cell expressing motility), we calculated the ratio of RFP expression-per-cell to YFP expression-per-cell,

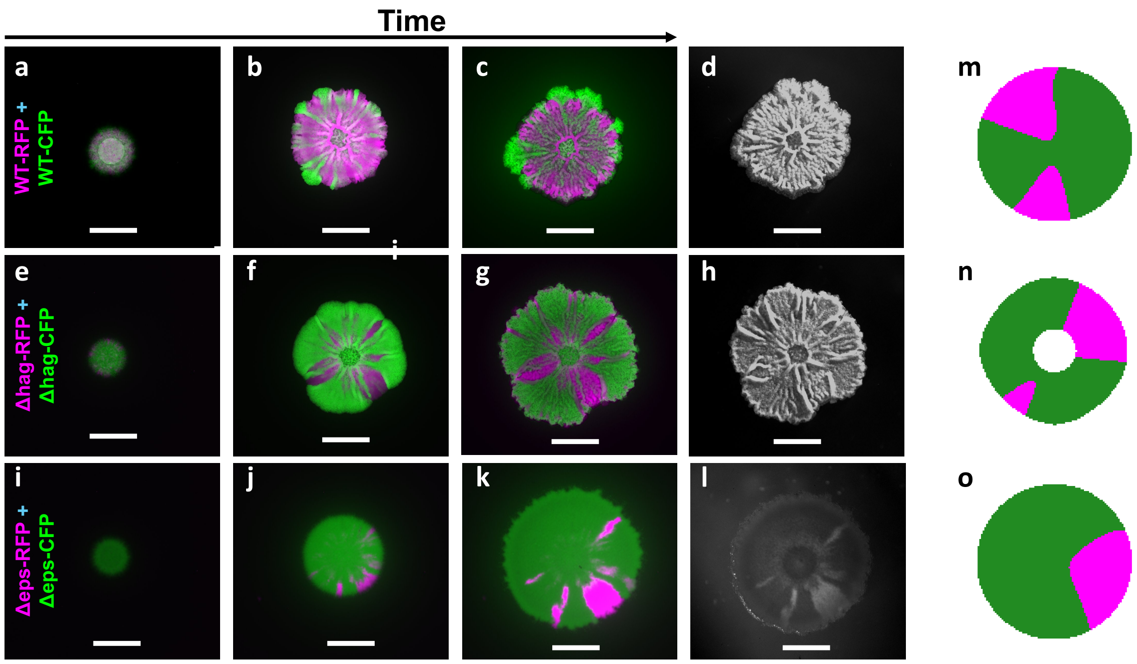


**Fig. S2** Competition experiments showing non-polar propagation patterns within growing biofilms initiated with isogenic strains carrying different fluorescent reporters: WT-RFP + WT-CFP (a–d), $\Delta$*hag*-RFP + $\Delta$*hag*-CFP (e–h), and $\Delta$*eps*-RFP + $\Delta$*eps*-CFP (i–l). For each competition, merged fluorescence images are shown for day 1 (a, e, i), day 3 (b, f, j), and day 6 (c, g, k), along with the corresponding bright-field image on day 6 (d, h, l). Scale bars represent 5000 $\mu m$. Corresponding simulation results of model 4 at 100 hour are shown in (m-o), for WT+WT, $\Delta$*hag* + $\Delta$*hag*, and $\Delta$*eps* + $\Delta$*eps*, respectively. Magenta and green represent isogenic strains with different fluorescent labels (green where strain 1 has fraction higher than 0.5 and magenta otherwise).

accounting for the different exposure times using:

$$\alpha_{\text{intrinsic}} = \frac{\overline{R} \cdot E_Y}{\overline{Y} \cdot E_R},$$

in which $\overline{R}$ is the mean RFP per cell, $\overline{Y}$ is the mean YFP per cell, $E_Y$ is the exposure time for the YFP channel, and $E_R$ is the exposure time for the RFP channel. For the cells analysed, we estimate $\alpha_{\text{intrinsic}} = 0.31$. This value was then multiplied by the ratio of exposure times used in the biofilm imaging, leading to $\alpha_{\text{exp}} = 0.42$, which is utilized as a known parameter $\alpha$ in the model.

We further calculated the ratio of mean YFP per cell and the average volume per cell to convert YFP intensity to volume fraction occupied by YFP expressing cells. After multiply this value by the ratio of exposure times used in biofilm experiment and single cell imaging, we estimate that under the exposure in biofilm experiment, the ratio of mean YFP per cell and the average volume per cell is 410. This value is used to initilize the fitting of parameter $\beta$ in the model.

##### 2.1.3 Comparison to biofilm measurements

To confirm that the values used in the calculation of $\alpha$ were reasonable, the single-cell images were compared to a number of biofilm images (example in fig. S11c). First, we plot the distribution of YFP and RFP pixel intensities (fig. S11a,b), noting significant probability mass at near zero intensity, corresponding to pixels with no YFP (or RFP) expressing cells, but possibly a small amount of auto-fluorescence. Considering only cellular contributions to the intensity by excluding all data below a threshold of 0.02, (Fig. S11d,e), we calculate a mean YFP (RFP) fluorescent intensity for pixels which contain YFP (RFP) -expressing cells.

Assuming the total fluorescent signal $S$ obtained from an object to be proportional to the number of fluorescent proteins $P$, the exposure time $E$, and the power of the objective $O$ (since this acts as a condenser for the excitation wavelengths), for some constant of proportionality $k$, we have

$$S = kPEO,$$

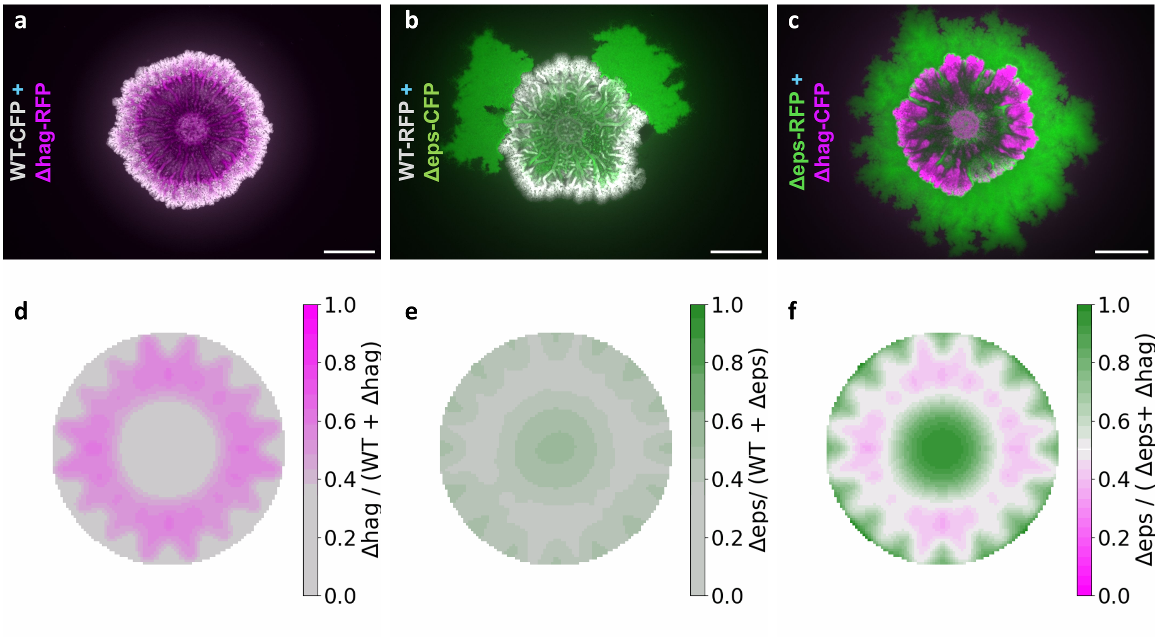


**Fig. S3** Final (day 7) merged biofilm images of mixed droplets of WT-CFP + Δ*hag*-RFP (a), WT-RFP + Δ*eps*-CFP (b), and Δ*eps*-RFP + Δ*hag*-CFP (c). Fluorescence images were captured using the respective filters, and false colors were applied to represent WT in gray, Δ*hag* in magenta, and Δ*eps* in green. Scale bars represent 5 mm. Corresponding simulation results of model 4 at 70 hours are shown in (d-f) for WT + Δ*hag*, WT + Δ*eps*, and Δ*eps* + Δ*hag*, respectively.

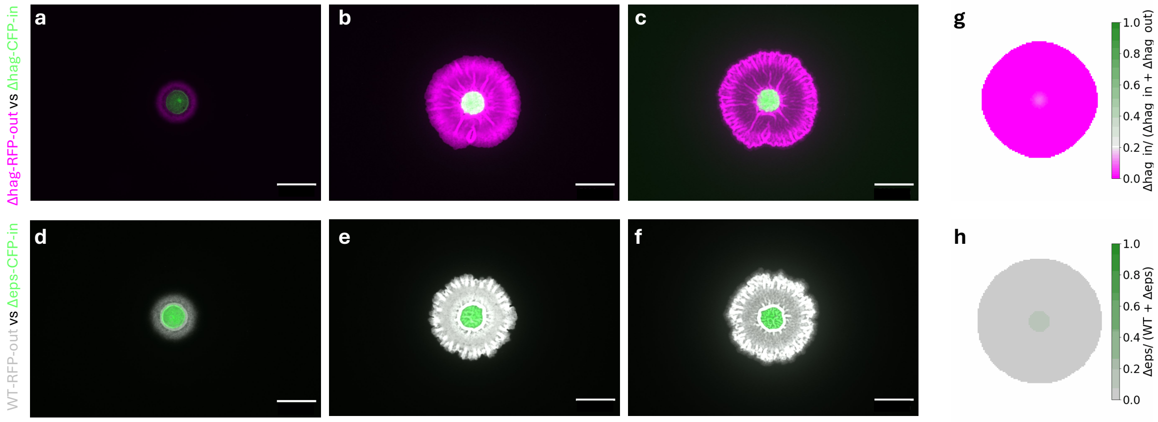


**Fig. S4** Additional bull's eye experiments to those shown in fig. 3, showing no escape over time for the combinations Δ*hag*-RFP-out-Δ*hag*-CFP-in (a-c) and WT-RFP-out-Δ*eps*-CFP-in (d-f). Panels correspond to day 1 (a, d), day 3 (b, e), and day 6 (c, f). Scale bars represent 5 mm. Corresponding simulation results of model 4 are shown in (g) and (h) for Δ*hag*-out-Δ*hag*-in at 50 hour, WT-out-Δ*eps*-in at 55 hour, respectively. The color bar in (g) is centered at 0.2 to help visualize the presence of Δ*hag*-in in the centre of the droplet.

or equivalently,

$$Pk = \frac{S}{EO}.$$

Assuming that the cells used for single-cell measurements have the same fluorescent properties as those in the biofilm (specifically that $Pk$ is conserved), we may estimate the number of cells that correspond to a single pixel (1.9 $\mu$m$^2$) of the biofilm $N_{\text{ppxl}}$ through

$$N_{\text{ppxl}} = \frac{P_{\text{cell}}k}{P_{\text{pxl}}k} = \frac{S_{\text{cell}} \cdot E_{\text{pxl}} \cdot O_{\text{pxl}}}{S_{\text{pxl}} \cdot E_{\text{cell}} \cdot O_{\text{cell}}}.$$

For the images obtained, we find $N_{\text{ppxl}} = 2.9$ cells per pixel for YFP-expressing cells, or $N_{\text{ppxl}} = 4.6$ cells per pixel for RFP-expressing cells. We compare this to the number of cells per pixel one

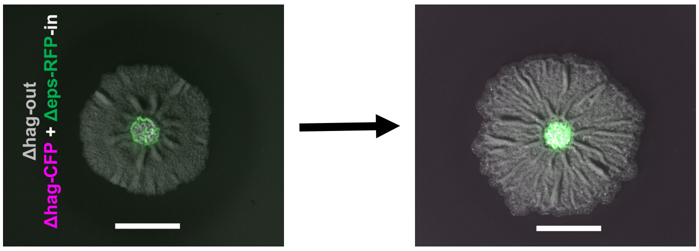


**Fig. S5** Bull's-eye experiments were conducted to assess the fate of an inner population consisting of a mixture of $\Delta$*hag*-CFP (magenta) and $\Delta$*eps*-RFP (green) cells, trapped within a $\Delta$*hag* (grey) biofilm. Images show day 2 (left) and day 6 (right) of biofilm development. No escape was observed in any of the 10 replicates. Scale bars represent 5000 $\mu$m.

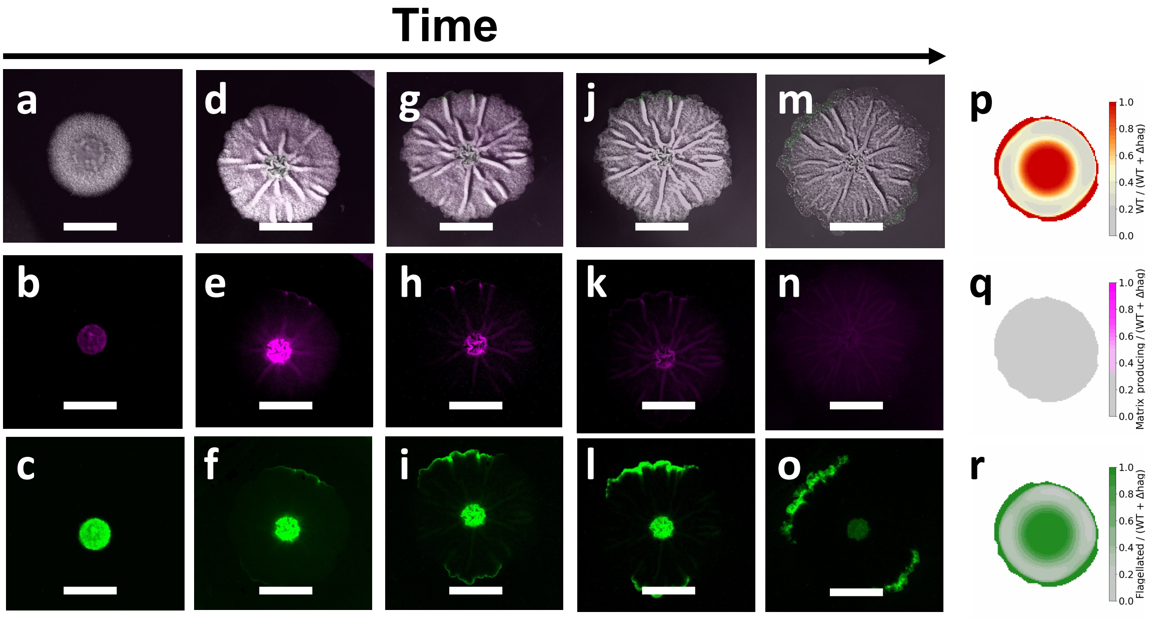


**Fig. S6** Tracking the gene expression throughout the escape of wild-type cells from a $\Delta$*hag* biofilm using the dual reporter for matrix and motility, ($\Delta$*hag*-out and dual-reporter-in). Images show bright-field (top row), matrix gene expression (magenta, middle row) and motility gene expression (green, bottom row) across an incremental timescale: Day 1 (a–c), Day 2 (d–f), Day 3 (g–i), Day 4 (j–l), and Day 7 (m–o). Corresponding simulation results of model 4 at 105 hour are shown in (p-r), where red identifies WT-in, magenta identifies the matrix cells within WT-in, and green identifies the flagellated cells within WT-in, respectively. Scale bars represent 5000 $\mu$m.

would expect at 100% packing fraction:

$$N_{\text{ppxl}} = \frac{\Delta x^2_{\text{biofilm}}}{A_{\text{cell}} \cdot \Delta x^2_{\text{cell}}},$$

in which $\Delta x_{\text{biofilm}}$ is absolute length per pixel in biofilm images (taken at a lower magnification), $\Delta x_{\text{cell}}$ is absolute length per pixel in single-cell images (taken at a higher magnification), and $A_{\text{cell}}$ is the mean single-cell area in pixels. Using this method, we find $N_{\text{ppxl}} = 0.46$ for both YFP and RFP-expressing cells. Taken together, these two measures of cell density imply that biofilm images have a cell volume fraction above 100% in a monolayer, and therefore that cells below the top biofilm layer (approximately $2.9/0.46 = 6.3$ or $4.6/0.46 = 10$ layers deep) contribute to the biofilm signal, which is expected in wide-field microscopy.

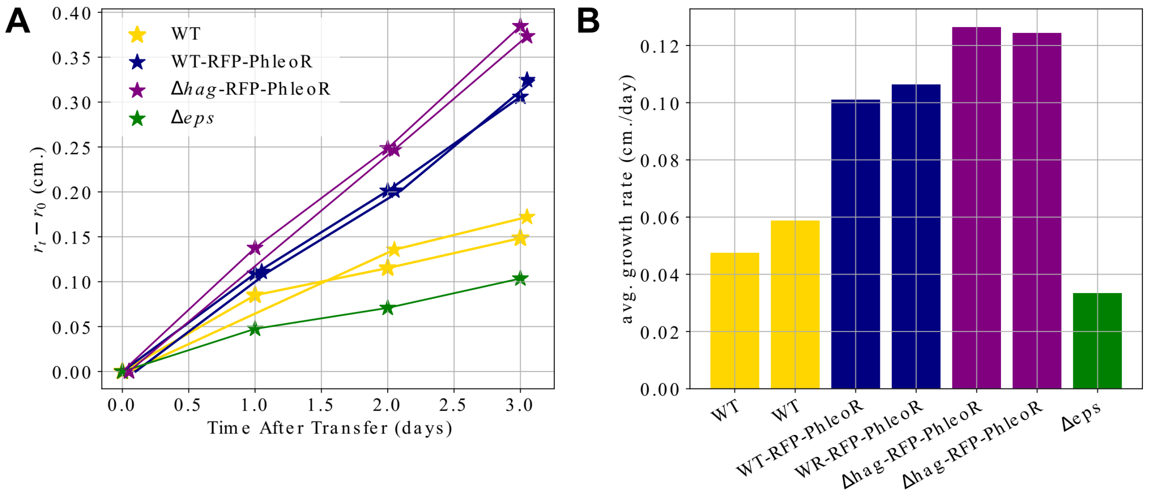


**Fig. S7** Growth of strains used in an experiment on 1.5% agar with MSgg over the course of three days after transfer to high concentration (10 $\mu$g/mL) of phleomycin. (a) The data plotted are colony radius subtracted by the radius at the time of transfer (t=0). For WT, WT-RFP-PhleoR, and $\Delta$*hag*-RFP-PhleoR, two replicates are shown with data points slightly jittered for clarity of overlapping data. Radius is estimated by calculating the area of binarised images of biofilms at each time point. (b) Average growth rate is calculated for each replicate by performing a linear fit of the data in (a).

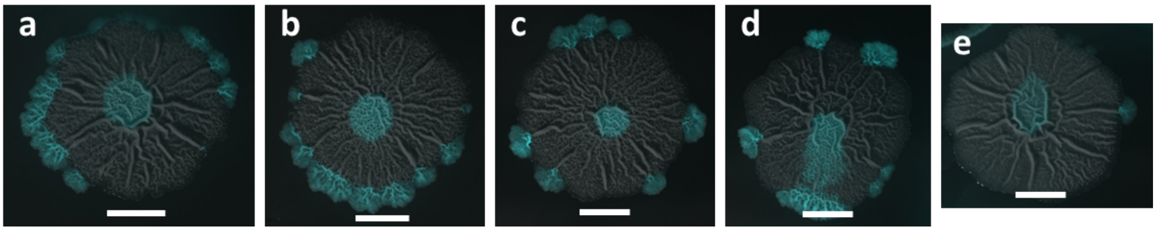


**Fig. S8** Biological replicates of successful escape of inner WT cells (cyan) on day 5 to the periphery of a WT biofilm (grey) after transfer to antibiotic plates at day 2. Scale bars represent 5000 $\mu$m.

### 3 Models of biofilm development

#### 3.1 Model variants

##### 3.1.1 Model 1: nutrient-dependent growth and stochastic switch

Model 1 is a reaction-diffusion model similar to a KPP-Fisher equation [3, 4] in which the flagellated and matrix cells grow according to Monod's law [5], with a nutrient-dependent maximum growth rate for matrix (and non-flagellated) cells and a constant maximum growth rate for flagellated (and non-matrix) cells. The switching rates for both flagellated to matrix and matrix to flagellated are small constants for WT and 0 for knock-out strains:

$$\begin{cases} \dfrac{\partial \phi_A}{\partial t} = D_A \nabla^2 \phi_A + \dfrac{k_{gA}\, n\, \phi_A}{n + K_A} - k_{dA}\phi_A - \mu_{AB}\phi_A + \mu_{BA}\phi_B, \\[2ex] \dfrac{\partial \phi_B}{\partial t} = D_B \nabla^2 \phi_B + \dfrac{(k_{gB0} + a\, n)\, n\, \phi_B}{n + K_B} - k_{dB}\phi_B + \mu_{AB}\phi_A - \mu_{BA}\phi_B, \\[2ex] \dfrac{\partial n}{\partial t} = D_n \nabla^2 n - \omega_1 \dfrac{k_{gA}\, n\, \phi_A}{n + K_A} - \omega_2 \dfrac{(k_{gB0} + a\, n)\, n\, \phi_B}{n + K_B}. \end{cases} \tag{1}$$

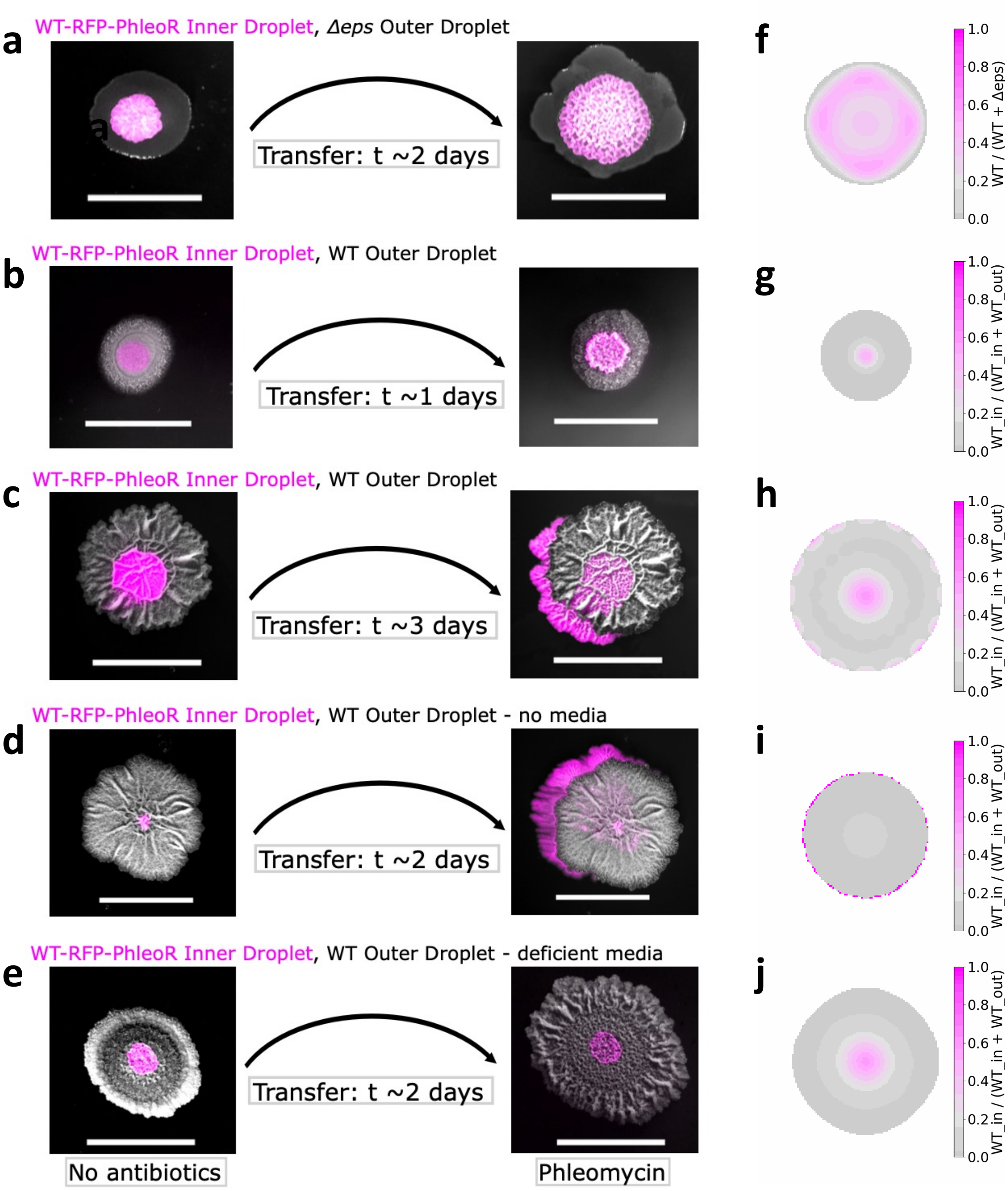


**Fig. S9** Additional control experiments with transfer on different media. The inner population is always shown in magenta and the outer one in grey. (a) $\Delta eps$-out-WT-RFP-PhleoR-in, no escape; (b) WT-RFP-PhleoR-in-WT-out transferred early (day 1), no escape; (c) WT-RFP-PhleoR-in-WT-out transferred late (day 3), escape; (d) WT-RFP-PhleoR-in-WT-out transferred on fresh plate with antibiotic and no media, escape; (e) WT-RFP-PhleoR-in-WT-out initially grown on media where all components are autoclaved, which prevents proper wrinkle formation, no escape. Scale bars represent 5 mm.(f-j) Corresponding simulation results from model 4, where transfer at day 1 corresponds to transfer at 20 hour, at day 2 corresponds to 40 hour and at day 3 corresponds to 45 hour. (f) and (g) are taken 5 hours after transfer, while all other snapshots are taken 25 hour after transfer process.

Here, $\phi_A$ is the flagellated cell volume fraction, $\phi_B$ is the matrix cell volume fraction, $n$ is the nutrient concentration, $\nabla^2$ is the Laplacian operator, $D_A, D_B, D_n$ are their respective diffusion coefficients, $k_{gA}$ and $k_{gB} = k_{gB0} + an$ are maximum per-capita growth rates ($a > 0$), $K_A, K_B$ are half-saturation constants, $k_{dA}$, $k_{dB}$ are death rates, $\mu_{AB}$, $\mu_{BA}$ are the phenotype switching rates, and $\omega_1$, $\omega_2$ are nutrient-yield coefficients. The values of the parameters are determined by fitting the radial fluorescent intensity profiles of the dual reporter *B. subtilis* strain as discussed in Sec. 3.3 (fig. S12, Table S3).

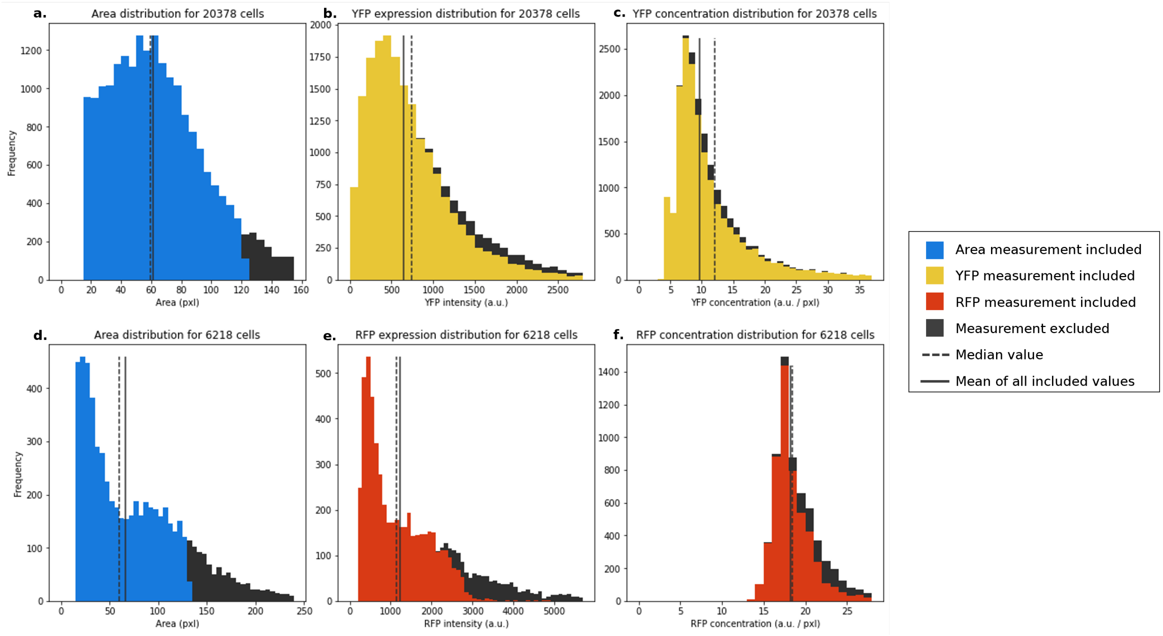


**Fig. S10** Histograms of cell properties obtained from single-cell agar pad measurements. **a.** Area of YFP-expressing cells. **b.** Total YFP-channel fluorescent intensity per YFP-expressing cell. **c.** Total YFP-channel fluorescent intensity per YFP-expressing cell, divided by cell area. **d.** Area of RFP-expressing cells. **e.** Total RFP-channel fluorescent intensity per RFP-expressing cell. **f.** Total RFP-channel fluorescent intensity per RFP-expressing cell, divided by cell area.

##### 3.1.2 Model 2: nutrient-dependent switch and constant maximum growth rate

As a counter-hypothesis to model 1, in model 2 we keep the maximal growth rates constant, and define the switching rates as nutrient-dependent, as summarized in the equations below:

$$\begin{cases} \dfrac{\partial \phi_A}{\partial t} = D_A \nabla^2 \phi_A + \dfrac{k_{gA}\, n\, \phi_A}{n + K_A} - k_{dA}\phi_A - (\mu_{AB0} + b\ n)\ \phi_A + \mu_{BA}\phi_B, \\ \dfrac{\partial \phi_B}{\partial t} = D_B \nabla^2 \phi_B + \dfrac{k_{gB}\, n\, \phi_B}{n + K_B} - k_{dB}\phi_B + (\mu_{AB0} + b\ n)\ \phi_A - \mu_{BA}\phi_B, \\ \dfrac{\partial n}{\partial t} = D_n \nabla^2 n - \omega_1 \dfrac{k_{gA}\, n\, \phi_A}{n + K_A} - \omega_2 \dfrac{k_{gB}\, n\, \phi_B}{n + K_B}. \end{cases} \tag{2}$$

Here, $k_{gA}, k_{gB}$, $\mu_{BA}$ are constants while $\mu_{AB} = \mu_{AB0} + bn$ $(b > 0)$.

After fitting with the dual reporter gene expression data (fig. S14, Table S3), this model can qualitatively reproduce the sequence of gene expression patterns in WT, but fails to capture the strain density profiles observed in mixed droplets of non-switching strains (fig. S13).

##### 3.1.3 Model 3: nutrient-dependent growth with enhanced diffusivity in wrinkles

Model 3 is identical to model 1 except for the wrinkles, which are defined as radial paths from the edge of the initial droplet to the edge of biofilm, and are activated after time $t_0 = 30\ h$. Within the wrinkles, flagellated cells have larger diffusion coefficient, so that $D_A \to D_A^{\text{eff}} = D_A\delta$, where $\delta$ is 1 before the wrinkles open, and increases to a higher value afterwards and within the wrinkles path (fig. S15, Table S3).

##### 3.1.4 Model 4: nutrient-dependent growth with wrinkling and cell transport

In order to explain the bull's eye experiments in fig. 3, we extend model 3 by taking into account the osmotic pressure generated by matrix production and the related fluid transport along the wrinkles.

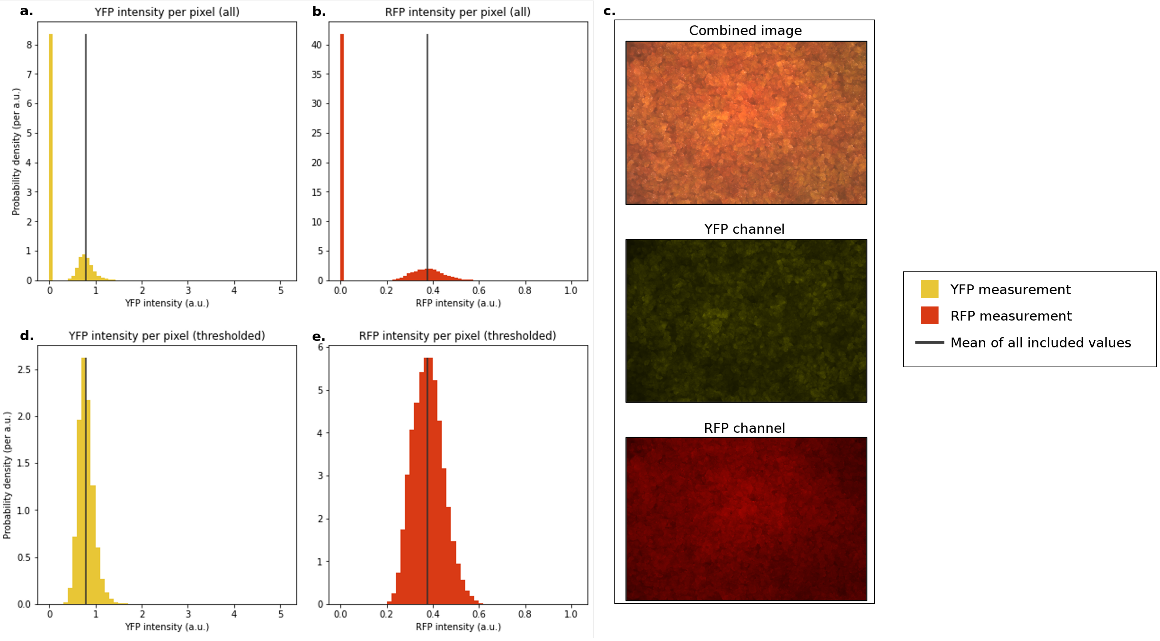


**Fig. S11** Histograms of fluorescent signal obtained from biofilm images. **a.** YFP signal per pixel. **b.** RFP signal per pixel. **c.** One of the biofilm images analyzed, showing the combined image (top) YFP channel (middle) and RFP channel (bottom). **d.** YFP signal per pixel after application of a minimum threshold. **e.** **RFP** signal per pixel after application of a minimum threshold.

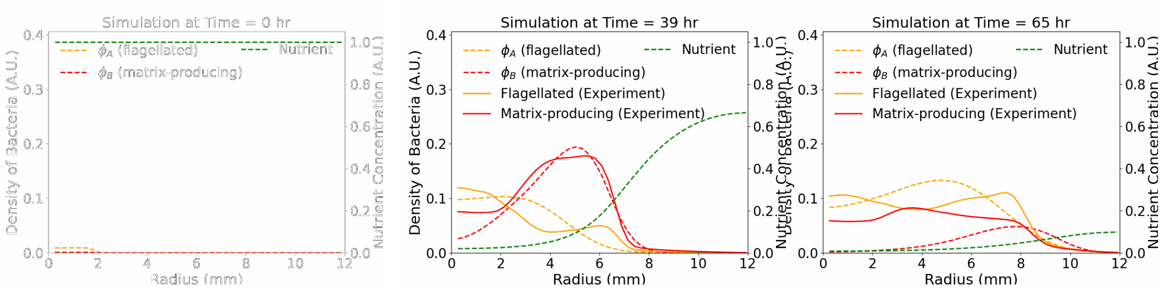


**Fig. S12** Comparison of gene expression profiles of the dual reporter biofilm between experiments (solid lines) and simulations of model 1 (dashed lines).

We define the volume fraction of a fluid (water) $\phi_f$ into our model, so that $\phi_A + \phi_B + \phi_f = 1$, under the simplifying assumption that the flagellated cells, the matrix cells and the fluid are present within the biofilm, while the nutrient field is inside the agar.

This leads to equations 3-6:

$$\frac{\partial \phi_A}{\partial t} = D_A \nabla^2 \phi_A + \frac{k_{gA}\, n\, \phi_A}{n + K_A} - k_{dA}\phi_A - \mu_{AB}\phi_A + \mu_{BA}\phi_B - \nabla \cdot (\phi_A \vec{v}_A)\, S \tag{3}$$

$$\frac{\partial \phi_B}{\partial t} = D_B \nabla^2 \phi_B + \frac{(k_{gB0} + an)\, n\, \phi_B}{n + K_B} - k_{dB}\phi_B + \mu_{AB}\phi_A - \mu_{BA}\phi_B - \nabla \cdot (\phi_B \vec{v}_B)\, S \tag{4}$$

$$\frac{\partial n}{\partial t} = D_n \nabla^2 n - \omega_1 \frac{k_{gA}\, n\, \phi_A}{n + K_A} - \omega_2 \frac{(k_{gB0} + an)\, n\, \phi_B}{n + K_B} \tag{5}$$

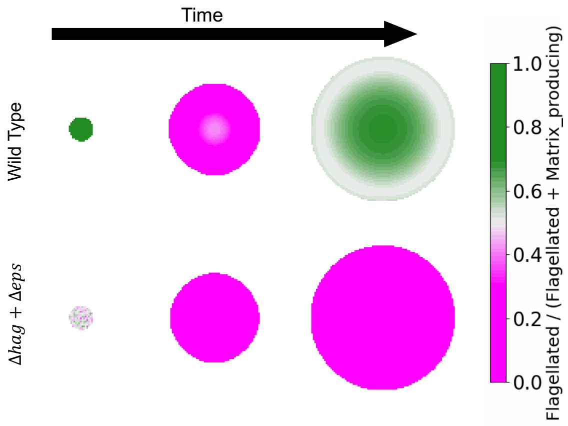


**Fig. S13** Gene expression pattern for wild-type (top) at 0 hour, 31 hour, 81 hour and for the mixture of knock-out strains (bottom) at 0 hour, 31 hour, 59 hour, based on model 2.

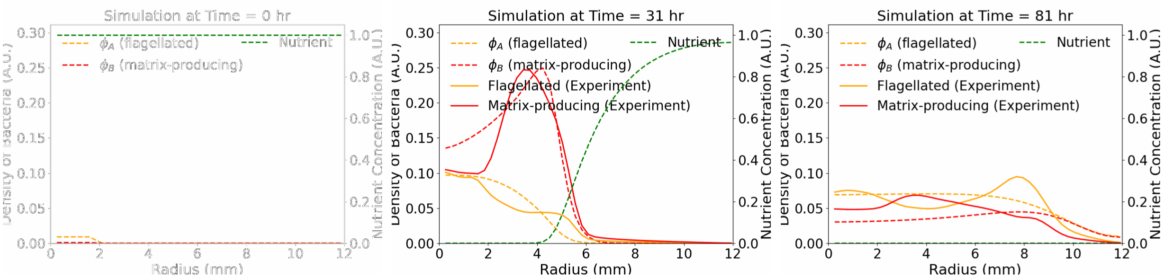


**Fig. S14** Comparison of gene expression profiles of the dual reporter biofilm between experiments (solid lines) and simulations of model 2 (dashed lines).

$$\begin{aligned}\frac{\partial \phi_f}{\partial t} = &-\left(D_A\nabla^2\phi_A + \frac{k_{gA}\, n\, \phi_A}{n+K_A} - k_{dA}\phi_A\right)\\ &-\left(D_B\nabla^2\phi_B + \frac{(k_{gB0}+an)\, n\, \phi_B}{n+K_B} - k_{dB}\phi_B\right) - \nabla\cdot(\phi_f\vec{v}_f)\, S.\end{aligned} \tag{6}$$

Here, $\vec{v}_A$, $\vec{v}_B$ and $\vec{v}_f$ represent the velocity of flagellated cells, matrix cells and fluid, respectively. $S = 1$ along the radial wrinkles after time $t_0$, and 0 otherwise.

Following the derivation in Ref. [6], we derive the equations of motion by minimizing the sum of free energy change and dissipation $R$, as defined in equation 7, with respect to $\vec{v}_A$, $\vec{v}_B$ and $\vec{v}_f$:

$$\begin{aligned}R = \int \mathrm{d}V \Bigg[&\frac{\eta_1}{2}|\vec{v}_A-\vec{v}_f|^2 + \frac{\eta_2}{2}|\vec{v}_A-\vec{v}_B|^2 + \frac{\eta_3}{2}|\vec{v}_B-\vec{v}_f|^2\\ &+\underline{\underline{\sigma}}_f\nabla\vec{v}_f + \underline{\underline{\sigma}}_A\nabla\vec{v}_A + \underline{\underline{\sigma}}_B\nabla\vec{v}_B + \frac{\partial f}{\partial\phi_A}\frac{\partial\phi_A}{\partial t} + \frac{\partial f}{\partial\phi_B}\frac{\partial\phi_B}{\partial t}\\ &- p\,\nabla\cdot\left(\phi_A\vec{v}_A + \phi_B\vec{v}_B + \phi_f\vec{v}_f\right)\Bigg],\end{aligned} \tag{7}$$

where $\eta_1$, $\eta_2$, $\eta_3$ are the friction coefficients between flagellated cells and fluid, flagellated cells and matrix cells, and matrix cells and fluid, respectively, $\underline{\underline{\sigma}}_f$, $\underline{\underline{\sigma}}_A$, $\underline{\underline{\sigma}}_B$ are the stress tensors of fluid, flagellated cells, and matrix cells, $f = f(\phi_A, \phi_B)$ is the mixing free energy, and $p$ (the pressure) is a Lagrange multiplier to enforce total volume conservation ($\nabla\cdot\left(\phi_A\vec{v}_A + \phi_B\vec{v}_B + \phi_f\vec{v}_f\right) = 0$). The time derivative of $\phi_A$ and $\phi_B$ are defined in equations 3 and 4. This yields equations 8-10:

$$\eta_1(\vec{v}_A-\vec{v}_f) + \eta_2(\vec{v}_A-\vec{v}_B) - \nabla\cdot\underline{\underline{\sigma}}_A + \phi_A\nabla\frac{\partial f}{\partial\phi_A} + \phi_A\nabla p = 0 \tag{8}$$

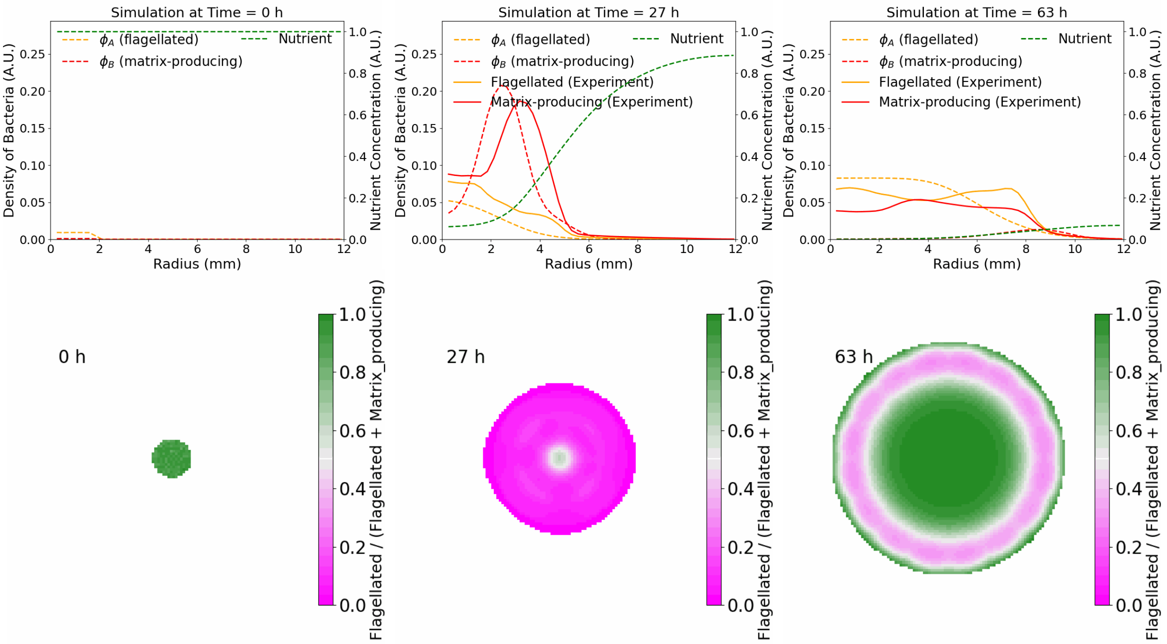


**Fig. S15** Top: comparison of gene expression profiles of the dual reporter biofilm between experiments (solid lines) and simulations of model 3 (dashed lines). Bottom: heatmap showing the proportion of flagellated (green) and matrix (magenta) cells throughout biofilm development in Model 3.

$$-\eta_2(\vec{v}_A - \vec{v}_B) + \eta_3(\vec{v}_B - \vec{v}_f) - \nabla \cdot \underline{\underline{\sigma}}_B + \phi_B \nabla \frac{\partial f}{\partial \phi_B} + \phi_B \nabla p = 0 \tag{9}$$

$$-\eta_1(\vec{v}_A - \vec{v}_f) - \eta_3(\vec{v}_B - \vec{v}_f) - \nabla \cdot \underline{\underline{\sigma}}_f + \phi_f \nabla p = 0. \tag{10}$$

We then add equations 8-10, which yields:

$$-\nabla \cdot \underline{\underline{\sigma}}_A - \nabla \cdot \underline{\underline{\sigma}}_B - \nabla \cdot \underline{\underline{\sigma}}_f + \phi_A \nabla \frac{\partial f}{\partial \phi_A} + \phi_B \nabla \frac{\partial f}{\partial \phi_B} + (\phi_A + \phi_B + \phi_f) \nabla p = 0. \tag{11}$$

We treat the system as a Newtonian fluid under quasi-static conditions, thus $\nabla \cdot \underline{\underline{\sigma}}_A = \lambda_A \nabla^2 \vec{v}_A$ where $\lambda_A$ is the viscosity of flagellated cells. We assume that non-flagellated/matrix cells negligibly flow throught the winkles, so $\vec{v}_B$ is set to 0 and thus $\nabla \cdot \underline{\underline{\sigma}}_B \approx 0$. The viscosity of water ($\lambda_f$) is expected to be much smaller than the viscosity of cells ($\lambda_A$ and $\lambda_B$) so also $\nabla \cdot \underline{\underline{\sigma}}_f \approx 0$. We define an osmotic pressure $\pi = \sum_i \phi_i \frac{\partial f}{\partial \phi_i} - f$, which leads to $\sum_i \phi_i \nabla \frac{\partial f}{\partial \phi_i} = \nabla \pi$. Plugging in $\phi_A + \phi_B + \phi_f = 1$, we then have:

$$\lambda_A \nabla^2 \vec{v}_A = \nabla p + \nabla \pi. \tag{12}$$

We set $\pi = \Pi \phi_B$, implying that only matrix cells drive osmotic pressure as they are the source of ECM, and find:

$$\lambda_A \nabla^2 \vec{v}_A = \nabla p + \Pi \, \nabla \phi_B. \tag{13}$$

If we momentarily consider the three-dimensional nature of a channel underneath a wrinkle, we expect that matrix cells ($\phi_B$) will predominantly be distributed along the walls of the channel, while the fluid ($\phi_f$) and flagellated cells ($\phi_A$) are likely mixed within the lumen of the channel. If we focus our analysis to the flow within the lumen of the channel (far away from the walls), then we can neglect the effect of matrix-fluid friction. As a result, equation 10 can be simplified to:

$$\eta(\vec{v}_f - \vec{v}_A) + \phi_f \nabla p = 0, \tag{14}$$

where we set $\eta_1 = \eta$, since it is the only remaining relevant frictional coefficient.

Model 4 then simplifies to:

$$\begin{cases} \dfrac{\partial \phi_A}{\partial t} = D_A \nabla^2 \phi_A + \dfrac{k_{gA}\, n\, \phi_A}{n + K_A} - k_{dA}\phi_A - \mu_{AB}\phi_A + \mu_{BA}\phi_B - \nabla \cdot (\phi_A \vec{v}_A)\, S, \\ \dfrac{\partial \phi_B}{\partial t} = D_B \nabla^2 \phi_B + \dfrac{(k_{gB0} + an)\, n\, \phi_B}{n + K_B} - k_{dB}\phi_B + \mu_{AB}\phi_A - \mu_{BA}\phi_B, \\ \dfrac{\partial n}{\partial t} = D_n \nabla^2 n - \omega_1 \dfrac{k_{gA}\, n\, \phi_A}{n + K_A} - \omega_2 \dfrac{(k_{gB0} + an)\, n\, \phi_B}{n + K_B}, \\ \dfrac{\partial \phi_f}{\partial t} = -\left( D_A \nabla^2 \phi_A + \dfrac{k_{gA}\, n\, \phi_A}{n + K_A} - k_{dA}\phi_A \right) \\ \qquad - \left( D_B \nabla^2 \phi_B + \dfrac{(k_{gB0} + an)\, n\, \phi_B}{n + K_B} - k_{dB}\phi_B \right) - \nabla \cdot (\phi_f \vec{v}_f)\, S, \\ \lambda_A \nabla^2 \vec{v}_A = \nabla p + \Pi\, \nabla \phi_B, \\ \eta\, (\vec{v}_f - \vec{v}_A) + \phi_f \nabla p = 0, \\ \nabla \cdot (\phi_A \vec{v}_A) = -\nabla \cdot (\phi_f \vec{v}_f), \\ \phi_A + \phi_B + \phi_f = 1. \end{cases} \tag{15}$$

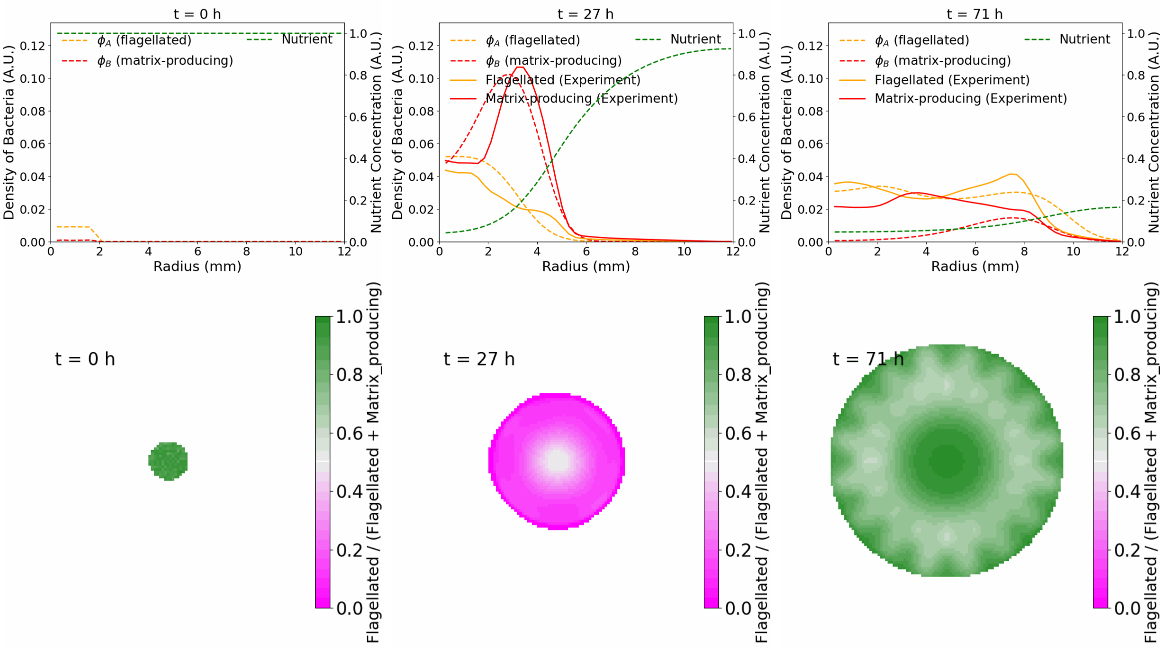


**Fig. S16** Top: comparison of gene expression profiles of the dual reporter biofilm between experiments (solid lines) and simulations of model 4 (dashed lines). Bottom: heatmap showing the proportion of flagellated (green) and matrix (magenta) cells throughout biofilm development in Model 4.

#### 3.2 Implementation of models

The numerical integration of the continuous models is written in Python 3. The models are solved either in polar coordinates (models 1 and 2 when the initial condition is polarly symmetric) or in Cartesian coordinates when polar symmetry is broken.

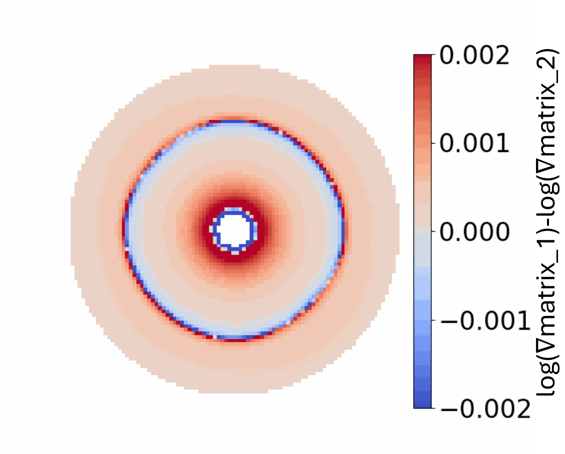


**Fig. S17** The heatmap of the difference in between the WT-in-Δhag-out setup and Δeps-in-Δhag-out setup. $\nabla\text{matrix}_1$ is the gradient of sum of volume fraction of all the matrix-producing cells in WT-in-Δhag-out setup, while $\nabla\text{matrix}_2$ is the gradient of sum of volume fraction of all the matrix-producing cells in Δeps-in-Δhag-out setup. Logarithmic function is utilized to make the difference clearer. Simulation is based on model 4. Screenshot is taken at 90 hour, which is around 15 hour before the screenshots in fig. 3(j-l) of main text.

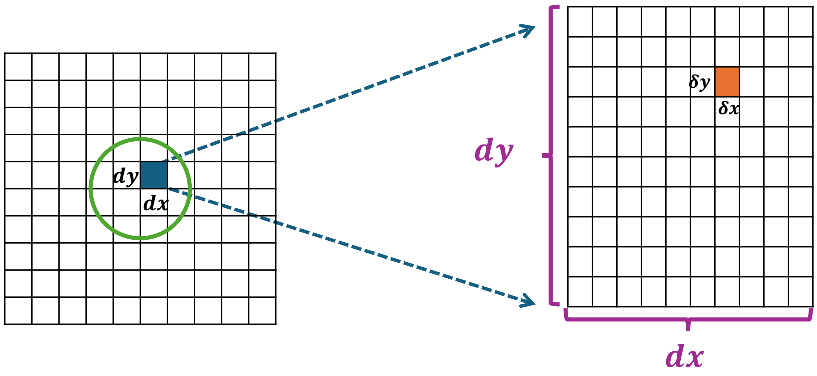


**Fig. S18** Schematic plot for initial condition. The green circle schematically shows the initial droplet. The big square on the left represent the grid of the simulation, while the right square zooms in the $dx \times dy$ from the left.

##### 3.2.1 Wild type and mixed droplets of non-flagellated and non-matrix strain

***Initial conditions***

All four models are implemented for the wild type bacteria and the mixture of non-flagellated and non-matrix strains.

For the polarly symmetric models 1 and 2, the simulations are initialized with a circular domain of radius $R = 2\text{mm}$ (approximately matching the size of the initial droplet in the experiments), $\phi_A + \phi_B = 0.01$ with $\phi_A/\phi_B = 10$, and $n(r, t = 0) = n_0 = 1$; boundaries are zero-flux.

For models 3 and 4, as the polar symmetry is broken, the initial condition follows a random process. As shown in fig. S18, we define for each position $(x, y)$ a bigger-block $dx \times dy$, each divided in a finer grid. Each small voxel in this finer grid, $\delta x \times \delta y$, is assigned either one of the two phenotypes. For the WT strain, the probability of being assigned a flagellated or matrix type are 0.9 and 0.1 respectively, and for the mixture of knock-outs is 0.5 each. We then count the proportion of flagellated voxels in the bigger block $dx \times dy$, and use this value divided by 100 as the initial value of $\phi_A$ in $(x, y)$. Similarly, we define $\phi_B$ counting the number of matrix voxels. To keep model 4 consistent, $\phi_f = 1 - \phi_A - \phi_B$. Boundaries are zero-flux.

.

***Simplification of $v_A$ calculation for model 4 integration***

To improve simulation efficiency without sacrificing the main ingredients of model 4, we initially focus on a wrinkle aligned along the $x$-axis, so that we can simplify the 2D problem to 1D. Our velocity field becomes

$$\vec{v} = (v_x(x,y,t), v_y(x,y,t)) = (v_x(x,t), 0) = (v(x,t), 0), \tag{16}$$

which neglects the $y$-component of the velocity as the wrinkles is predominatly extended along the $x$-direction and any $y$-dependence of the $x-$component. By replacing equation 13 into equation 14, we then find

$$\eta(v_f - v_A) + \lambda_A \phi_f v_A'' - \Pi \phi_f \phi_B' = 0, \tag{17}$$

where the derivatives are all taken with respect to $x$. In addition, equation $\nabla \cdot (\phi_A \vec{v}_A + \phi_f \vec{v}_f) = 0$ simplifies to $\frac{\partial}{\partial x}(\phi_A v_A + \phi_f v_f) = 0$. Thus, $\phi_A v_A + \phi_f v_f = C$ where $C$ is a constant, leading to

$$\frac{\eta C}{\phi_f} - \eta \left( \frac{\phi_A}{\phi_f} + 1 \right) v_A + \lambda_A \phi_f v_A'' - \Pi \phi_f \phi_B' = 0. \tag{18}$$

$C$ is estimated in the quick search round of model 4, yielding $C = 0$.

The simplification holds for a wrinkle running along the $x$-axis. Assuming that radial wrinkles at different angles are identical to each other, we apply the same 1D equations to all the wrinkles along their direction of propagation. For each time step of the simulations in model 4, $v_A$ is solved for the current values of $\phi_A$, $\phi_B$ and $\phi_f$, and then updated at each time step.

***Solve the equations and visualize the results***

The wrinkles open at time $t = 30h$ and are characterized by a half-width 0.275 mm. We place 16 wrinkles equally distributed around the biofilm. The wrinkle propagate from the edge of the initial droplet the up to the edge of the biofilm, defined as when $\phi_A + \phi_B$ drops below $10^{-4}$.

In all models, the system of equations is solved with the python package *solve_ivp* and method $RK45$ (Runge–Kutta methods [7, 8]). The calculation of $v_A$ at each time step is done using the python package *spsolve*.

The visible threshold for the biofilm in the figures is set at $5 \cdot 10^{-5}$ for each phenotype. The simulation box is 12 mm wide, with $\Delta x = 0.265$. The simulations are run for 90 hours. The time step $dt$ is automatically self-adjusted to balance the speed and accuracy, while the data is output every 0.667 hour in line with the experimental frame rate.

##### 3.2.2 Mixed droplets

Model 4 is implemented for the simulation of mixed droplets. For the simulation of mixed droplets with more than one strains (figs. S2 and S3), the initial ratio is 1:1, and the initial total cell volume fraction is 0.01, i.e., $1 - \phi_f = 0.01$. When the WT is present in the mixed droplet, the flagellated-matrix ratio in WT strain is 10:1. The simulations are run until 100 hours for mixed droplet of isogenic strains with different fluorescent reporters, and 70 hours for other mixed droplets. All other simulation-related setup are the same as in Section 3.2.1.

##### 3.2.3 Bull's eye simulations - no transfer

Model 1, 3, 4 are implemented for bull's eye (no transfer) initial conditions. The initial size is 1 mm for the inside strain and 3 mm for the outside strain. For droplets with more than one phenotypes, the initial state follows the same random initial configuration as described in Section 3.2.1. For droplets with only one phenotype, the initial volume fraction for that phenotype is 0.01. The simulation time is 65 hour for model 1, 60 hour for model 3, and 105 hour for model 4. All other setups are the same as described in 3.2.1 for the corresponding model. Simulation results are shown in fig. 3, S4, S6 and S17.

##### 3.2.4 Bull's eye simulations - with transfer

Model 4 is implemented for the bull's eye transfer experiments. The initial setup is the same as in the bull's eye experiment with no transfer considered (Section 3.2.3). The transfer process occurs at 40 hours after inoculation (unless stated otherwise). After transfer, the nutrient is reinitialized to the initial value, and the transport ability, growth rate, switch rate, and diffusion coefficient of the outside strain are decreased to 50% of the original value, in line with the experimental results in fig. S7. We run the simulations until 25 hours after the transfer process.

In these conditions, we set the visible threshold as stated earlier, with the exception of the case in which no fresh nutrients are added after the transfer (fig. S9i). Here, we set the visible threshold to $10^{-6}$ to accommodate the overall lower cell abundance in this condition.

#### 3.3 Gene expression data analysis and parameterization of the models

##### 3.3.1 Data analysis of gene expression profiles

The average values of the flagellated cell intensity (YFP signal) and matrix cell intensity (RFP signal) from the dual reporter time-lapse experiments at each radius of the biofilm are extracted using Fiji (Version 1.54p) [9]. The background intensity is removed by subtracting the intensity values outside the edge of the biofilm and the remaining negative values (close to 0 near the edge of the biofilm) are set to 0. The RFP signal is first normalized by $\alpha$ to account for the different intensities between RFP and YFP, as discussed in Section 2.1.2. Then, both the YFP signal and the normilized RFP signal are processed via a Gaussian filter with $\sigma = 10$.

##### 3.3.2 Model parameterization by fitting gene expression data

In order to obtain the parameter values of our 4 models, we fit the time-dependent volume fraction simulation profiles from our models with the RFP and YFP signals processed as described above. The parameter $\beta$ is introduced to link the volume fraction with the fluorescence intensity (See Section 2.1.2).

We calculate the root mean square error (RMSE) between the experimental data and simulation profiles for RFP and YFP separately. To achieve a small relative error in both YFP and RFP fitting procedures, we define a weight parameter as

$$w = \frac{\max(\text{RFP})}{\max(\text{YFP})} \tag{19}$$

describing the ratio of the maximum value of processed RFP data and YFP data. Then we select the bigger value between the $w * \text{RMSE(YFP)}$ and RMSE(RFP), i.e.,

$$\overline{\text{RMSE}} = \max(w * \text{RMSE(YFP)}, \text{RMSE(RFP)}). \tag{20}$$

We then used the Nelder–Mead method [10] with initial simplex scaling factor 0.05 and total number of iterations equal to 1000 to find the parameter sets leading to the smallest $\overline{\text{RMSE}}$ .

For model 1 and 2, which display polar symmetry, the simulation profiles are directly compared to the average radial intensity collected in the experiments. For model 3 and 4, wrinkles break the polar symmetry. To improve the efficiency of the fitting procedure, we start from a simpler version of model 3 and 4 (model 3* and 4*) in which the cells can disperse out everywhere in all directions as if everything is a large wrinkle. The models 3* and 4* allow us to use a polarly symmetric configuration to fit the parameters faster and reach a parameter set that is close to our best-fit result for models 3 and 4. We refer to this parameter set as the preliminary best fit. This preliminary best fit is then used as a starting point to fit the parameters in Cartesian coordinate for models 3 and 4. For each iteration, after calculating the volume fractions at each $(x, y)$ point, the volume fractions at the same radius are averaged and fit against the experimental data. The fitting process runs for another 50 iterations to find the final best-fit in Cartesian coordinate.

Table S3 summarizes the final parameters for each model, and comparison of experimental and simulation gene expression profiles after the fitting procedure are shown in fig. S12, S14-S16 for the 4 different models.

### 4 Cell dynamics within the wrinkle network

One of the key assumptions of model 4 is that, while both matrix producing and flagellated cells contribute to building the osmotic pressure necessary to draw liquid inwards (and push cells outwards),

**Table S3** Parameters of the four models found by fitting the experimental gene expression data from the dual reporter

| Symbol | Description | Model 1 | Model 2 | Model 3 | Model 4 | Unit |
|---|---|---|---|---|---|---|
| $D_n$ | Nutrient diffusion coefficient | 0.91 | 0.50 | 1.0 | 0.87 | $mm^2\,h^{-1}$ |
| $k_{gA}$ | Maximum growth rate (flagellated cells) | 0.26 | 0.15 | 0.13 | 0.25 | $h^{-1}$ |
| $K_A$ | Half–saturation constant (flagellated cells) | 0.034 | 0.058 | 0.010 | 0.14 | $A.U.^{-1}$ |
| $D_A$ | Diffusion coefficient (flagellated cells) | 0.040 | 0.039 | 0.057 | 0.032 | $mm^2\,h^{-1}$ |
| $k_{gB}$ | Maximum growth rate (matrix cells) | 0.092 | 0.35 | 0.11 | 0.10 | $h^{-1}$ |
| $K_B$ | Half–saturation constant (matrix cells) | 0.10 | 0.020 | 0.11 | 0.011 | $A.U.^{-1}$ |
| $D_B$ | Diffusion coefficient (matrix cells) | 0.0082 | 0.022 | 0.0049 | 0.016 | $mm^2\,h^{-1}$ |
| $k_{dA}$ | Death rate (flagellated cells) | 0.085 | 0.0075 | 0.028 | 0.068 | $h^{-1}$ |
| $k_{dB}$ | Death rate (matrix cells) | 0.13 | 0.035 | 0.19 | 0.24 | $h^{-1}$ |
| $\omega_1$ | Yield coefficient (flagellated cells) | 0.25 | 0.62 | 1.33 | 0.52 | A.U. |
| $\omega_2$ | Yield coefficient (matrix cells) | 2.78 | 4.94 | 3.40 | 5.10 | A.U. |
| $b$ | Nutrient–dependent switch rate factor | — | 0.89 | — | — | $A.U.^{-1}\,h^{-1}$ |
| $a$ | Nutrient–dependent growth rate factor | 1.35 | — | 1.41 | 0.76 | $A.U.^{-1}\,h^{-1}$ |
| $\mu_{AB}$ | Switch rate (flagellated to matrix) | $1.6\times 10^{-5}$ | 0.019 | $3.1\times 10^{-4}$ | $1.7\times 10^{-4}$ | $h^{-1}$ |
| $\mu_{BA}$ | Switch rate (matrix to flagellated) | $5.9\times 10^{-4}$ | 0.029 | $5.6\times 10^{-4}$ | $2.7\times 10^{-5}$ | $h^{-1}$ |
| $\delta$ | Diffusion multiplier within wrinkles | — | — | 10.34 | — | - |
| $\beta$ | YFP per cell to cell volume fraction ratio | 187.62 | 198.16 | 281.06 | 513.68 | A.U. |
| $\lambda_A$ | Viscosity of flagellated cells | — | — | — | 0.58 | Pa h |
| $\eta$ | Friction between flagellated cells and fluid | — | — | — | 0.82 | Pa h $mm^{-2}$ |
| $\Pi$ | Osmotic compressibility | — | — | — | 14.93 | Pa |

*Notes.* For nutrient–dependent switching, $\mu_{AB}$ shown here corresponds to the basal switch rate (i.e., $n = 0$). For nutrient–dependent growth, $k_{gB}$ denotes the growth rate when $n = 0$.
A.U. means arbitrary unit.

only flagellated cells can effectively flow through the wrinkles ($\vec{v_B} \ll \vec{v_A}$). Testing this assumption within the standard development of the biofilm is technically challenging, as it is difficult to identify and track the relatively few cells that move inside the wrinkle within the much larger ensemble of cells that forms the wrinkle. To bypass this problem and provide a quantification of how flagellated vs. non-flagellated cells explore the wrinkle network of the biofilm, we follow the protocol and data analysys below.

#### 4.1 Introduction and imaging of cells in biofilms channels

Wild-type NCIB 3610 *B. subtilis* biofilms were grown on MSgg agar for 2 days as described earlier. Overnight cultures of WT-RFP and Δhag-CFP in LB were concentrated 10 times through centrifugal precipitation and redisolving. Concentrated overnight cultures were mixed in equal proportions. A small hole was gently made near the central region of the biofilm at the origin of a large wrinkle with a P10 pipette tip. A 5 $\mu$L PBS buffer was injected into the opening to wet the wrinkle and reduce capillary effects before the addition of 2 $\mu$L cell mixture. Immediately ($\sim$ 20 seconds) after, the sample was imaged from below, through the agar, using fluorescent timelapse microscopy as described below, with 24-second intervals between frames. Images were taken with a 2.3x objective. A representative time-series of the cell spatial distribution is shown in fig. S19b-f, where the yellow arrow points out a region of the wrinkle network that is readily explored by the WT cells (green), but not by the non-flagellated Δ*hag* cells (magenta).

A control photobleaching experiment was performed to assess the effect of fluorescence imaging on *Bacillus* cells during the injection assay fig. S20. A similar concentrated cell mixture was placed on a glass slide and covered with a coverslip; a coverslip sealant was applied to create a closed environment suitable for time-lapse imaging. Identical imaging parameters were used to capture images at a similar resolution. Six $\tilde{3}0 \times 30$ $\mu$m regions of interest were selected to track CFP and RFP fluorescence intensity over time. Box F represents background noise.

#### 4.2 Data analysis and statistics

To better quantify the time scales associated with the dynamics of the flagellated and non-flagellated cells across the biofilm, we divide the biofilm images in 62x62 tiles and track the average RFP and CFP fluorescent signal change with respect to time for each grid position (example of profile shown in fig. S19g).

Firstly, we apply a moving average of the time series data with a window size of the 13.30 minutes to smooth out fluctuations. We then select those curves which maxima-minima difference is greater

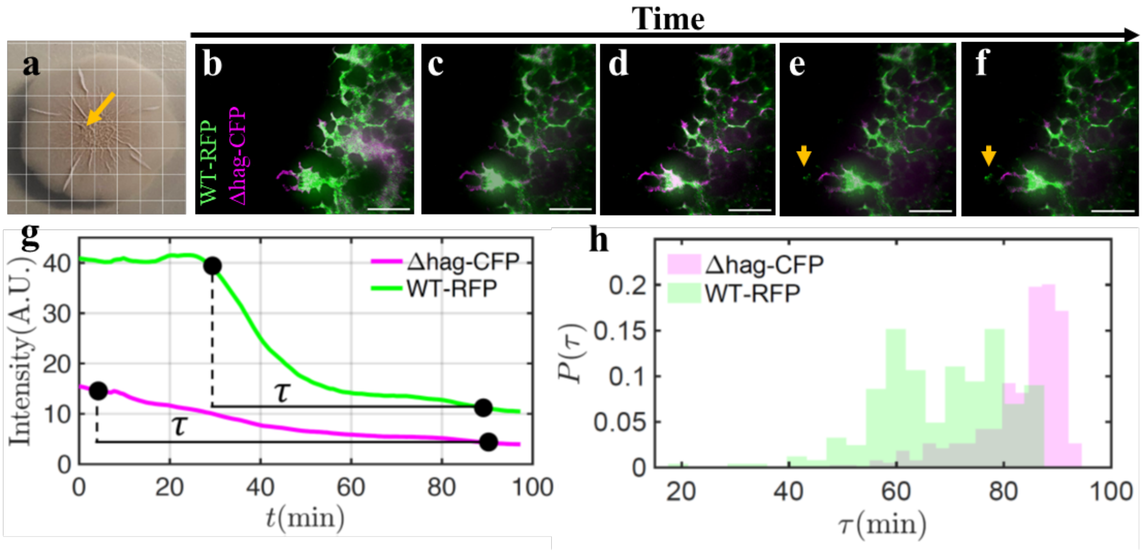


**Fig. S19** Dynamics within biofilm wrinkles reveal significant movement of WT cells, reaching regions inaccessible to non-flagellated Δ*hag*-CFP cells. (a) Two-day-old biofilm used for the tracking study; the yellow arrow indicates the injection site. (b–f) Time-series images showing the movement of Δ*hag*-CFP and WT-RFP cells within biofilm wrinkles. Fluorescent images were generated by merging RFP and CFP channel captures. Yellow arrows in panels (e) and (f) highlight the appearance of green sectors in previously empty regions, indicating the movement of WT-RFP (green) cells into areas that were inaccessible to non-flagellated Δ*hag*-CFP (magenta) cells. Plot of fluorescent intensity versus time for Δ*hag*-CFP and WT-RFP as shown in (g). The relaxation time ($\tau$) is defined as the time taken for intensity to decrease from 95% of its maximum value to 105% of its minimum value. In (h), the probability distribution function $P(\tau)$ of relaxation time ($\tau$) is shown for both the Δ*hag*-CFP and WT-RFP. Scale bars represent 500 $\mu$m.

than or equal to $g_{\mathrm{cut}} = 10$. The qualitative results remain consistent for $g_{\mathrm{cut}} = 2, 4, 6$, and 8. Choosing the higher cutoff filter out curves that only fluctuate around the mean without showing clear growth or decay; therefore we use $g_{\mathrm{cut}} = 10$ for our analysis. For non-flagellated cells, the fluorescence curves mostly decay monotonically, as shown in the representative plot in fig. S19g). We define the relaxation time as the time taken for a curve to decay from its 95% of its maximum values to 105% of its minimum value. Almost all curves (379) of the non-flagellated cells decreases monotonically, whereas a significant fraction of flagellated cell curves (131) increases monotonically and few curves (27) first increase and then decrease . Representative plots of the monotonically increasing curve and the "increase-then-decrease" curve are shown in the fig. S21.

The normalized histogram of the decay timescales for the different strains across the whole biofilm is reported in fig. S19h. The analysis clearly shows that non-flagellated cells (Δ*hag*) are characterized by significant longer timescales compared to the wild-type, which can be interpreted as a lower overall flux across different regions of the biofilms. Interestingly, the WT cells display a bimodal distribution. The origin of this bimodality is unclear, but a potential explanation could be found in the inherent phenotypic heterogeneity of the WT cells, since they can randomly switch from flagellated to matrix state.

It is important to note that cell movement and transport across the network in these conditions are driven by the sudden introduction of a large number of cells, so a different mechanism from what we expect during biofilm development, and what our model attempts to describe mathematically. Nevertheless, these results support the idea that non-flagellated cells are less capable of moving across the network compared to their flagellated counterparts. Whether this is due to higher rates of attachment to the walls of the wrinkles or limited swimming ability remains to be determined.

### References

[1] Asally, M. *et al.* Localized cell death focuses mechanical forces during 3d patterning in a biofilm. *Proceedings of the National Academy of Sciences* **109**, 18891–18896 (2012). URL https://www.pnas.org/doi/abs/10.1073/pnas.1212429109.

[2] Stringer, C., Wang, T., Michaelos, M. & Pachitariu, M. Cellpose: a generalist algorithm for cellular segmentation. *Nature methods* **18**, 100–106 (2021).

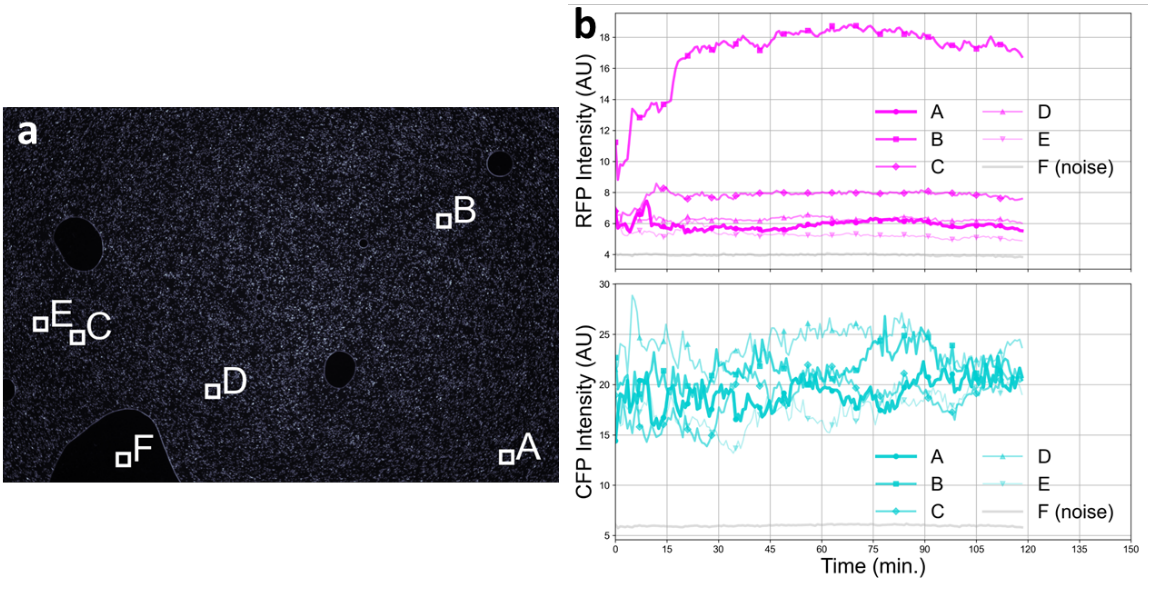


**Fig. S20** Cell mixtures in the channel experiment were imaged under consistent conditions to assess photobleaching effects on fluorescence. (a) Representative timelapse snapshot showing regions A–F, each $\tilde{3}0 \times 30$ $\mu$m. Region F contained no cells and served as a noise control. (b) Average CFP and RFP intensities for regions A–F over time (min) were quantified and plotted.

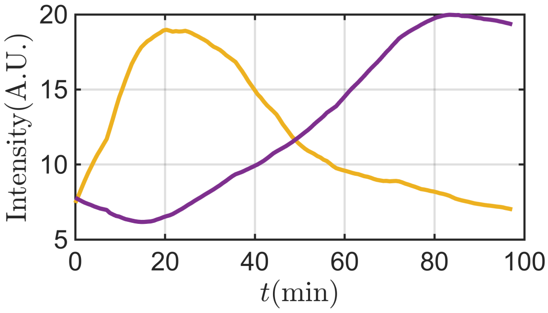


**Fig. S21** Representative plots of fluorescence intensity versus time for WT-RFP within the channel, analogous to fig. S19g), showing both growth–decay dynamics and purely growing dynamics.

[3] Fisher, R. A. The wave of advance of advantageous genes. *Annals of eugenics* **7**, 355–369 (1937).

[4] Kolmogoroff, A., Petrovsky, I. & Piscounoff, N. in *Study of the Diffusion Equation with Growth of the Quantity of Matter and its Application to a Biology Problem* (ed.Pelcé, P.) *Dynamics of Curved Fronts* 105–130 (Academic Press, San Diego, 1988). URL https://www.sciencedirect.com/science/article/pii/B9780080925233500149.

[5] Monod, J. The growth of bacterial cultures. *Selected Papers in Molecular Biology by Jacques Monod* **139**, 606 (2012).

[6] Seminara, A. *et al.* Osmotic spreading of bacillus subtilis biofilms driven by an extracellular matrix. *Proceedings of the National Academy of Sciences* **109**, 1116–1121 (2012).

[7] Runge, C. Über die numerische auflösung von differentialgleichungen. *Mathematische Annalen* **46**, 167–178 (1895).

[8] Kutta, W. *Beitrag zur näherungsweisen Integration totaler Differentialgleichungen* (Teubner, 1901).

[9] Schindelin, J. *et al.* Fiji: an open-source platform for biological-image analysis. *Nature methods* **9**, 676–682 (2012).

[10] Nelder, J. A. & Mead, R. A simplex method for function minimization. *The computer journal* **7**, 308–313 (1965).
